# Massively parallel genetic perturbation reveals the energetic architecture of an amyloid beta nucleation reaction

**DOI:** 10.1101/2024.07.24.604935

**Authors:** Anna Arutyunyan, Mireia Seuma, Andre J. Faure, Benedetta Bolognesi, Ben Lehner

**Affiliations:** Wellcome Sanger Institute, Cambridge, UK; Institute for Bioengineering of Catalonia (IBEC), The Barcelona Institute of Science and Technology (BIST), Baldiri Reixac 10-12, 08028, Barcelona, Spain; Centre for Genomic Regulation (CRG), The Barcelona Institute for Science and Technology (BIST), Barcelona, Spain; Universitat Pompeu Fabra (UPF), Barcelona, Spain; Institució Catalana de Recerca i Estudis Avançats (ICREA), Barcelona, Spain; Current address: ALLOX, C/ Dr. Aiguader, 88, PRBB Building, 08003 Barcelona, Spain

## Abstract

Amyloid protein aggregates are pathological hallmarks of more than fifty human diseases but how soluble proteins nucleate to form amyloids is poorly understood. Here we use combinatorial mutagenesis, a kinetic selection assay, and machine learning to massively perturb the energetics of the nucleation reaction of amyloid beta (Aβ42), the protein that aggregates in Alzheimer’s disease. In total, we quantify the nucleation rates of >140,000 variants of Aβ42. This allows us to accurately quantify the changes in reaction activation energy for all possible amino acid substitutions in a protein for the first time and, in addition, to quantify >600 energetic interactions between mutations. The data reveal the simple and interpretable genetic architecture of an amyloid nucleation reaction. Strikingly, strong energetic couplings are rare and identify a subset of structural contacts in mature fibrils. Together with the activation energy changes, this strongly suggests that the Aβ42 nucleation reaction transition state is structured in a short C-terminal region, providing a structural model for the reaction that may initiate Alzheimer’s disease. We believe this approach can be widely applied to probe the energetics and transition state structures of protein reactions.

## Introduction

Amyloid fibrils are supramolecular fibrous protein assemblies in which β-strands are stacked along the long axis of each fibril in an ordered ‘cross-β’ structure^1^. Specific amyloid assemblies define most human neurodegenerative diseases, including Alzheimer’s disease, Parkinson’s disease and frontotemporal dementia. Fibrils of amyloid beta (Aβ42) are, for example, a pathological hallmark of Alzheimer’s disease and variants in Aβ42 cause familial Alzheimer’s disease^2,3^. In total, at least 50 human disorders are associated with the formation of amyloid fibrils of more than 30 different proteins^4^. Beyond human pathology, amyloids are present in all kingdoms of life and possess functions in a wide variety of organisms, including humans^5^.

Mature amyloid fibrils are stable structures and normally irreversible states. For many proteins they are likely to be the thermodynamically most favoured state at high protein concentration^6^. The conversion of soluble proteins to amyloid fibrils occurs through nucleation-and-growth processes, with nucleation being the rate-limiting step^7^. Most proteins do not form amyloids under physiological conditions because of kinetic control: the high free energy barrier of the nucleation reaction means fibrils never form on human timescales^8,9^. Understanding why a small subset of proteins aggregate in human diseases therefore requires understanding nucleation reactions. Nucleation is also the key process to prevent in order to stop the formation and spreading of amyloids. Preventing nucleation would stop both the formation of mature fibrils and the production of potentially toxic fibril intermediates and oligomeric species produced during fibril assembly^10^. Nucleation reactions are therefore the key processes to understand and prevent in order to stop amyloid formation and amyloid-associated toxicity in human diseases.

Thanks primarily to developments in cryogenic electron microscopy (cryo-EM), the atomic structures of many amyloid fibrils have now been determined^11^, including fibrils extracted from human brains^12^. These structures have revealed that proteins can adopt different filament structures (fibril polymorphs) with, at least in some cases, different amyloid folds associated with different clinical conditions^11^. Time-resolved cryo-electron microscopy has also been used to characterise the *in vitro* assembly of amyloids, revealing a diversity of folds that appear and disappear as fibrillation proceeds^13,14^. Protein folding intermediates implicated in the initiation of amyloid formation have also been partially characterised by NMR, revealing native-like structures^15,16^.

The rate of a nucleation reaction depends on the difference in energy (the activation energy, E_a_) between the highest energy state along the reaction coordinate (the transition state) and the initial soluble state (Fig. 1a). Due to their high energy and transient nature, transition states are notoriously difficult to characterise. In contrast to folding intermediates^13–16^, transition states have therefore not been structurally characterised for any amyloid nucleation reaction^17^.

**Figure 1.**
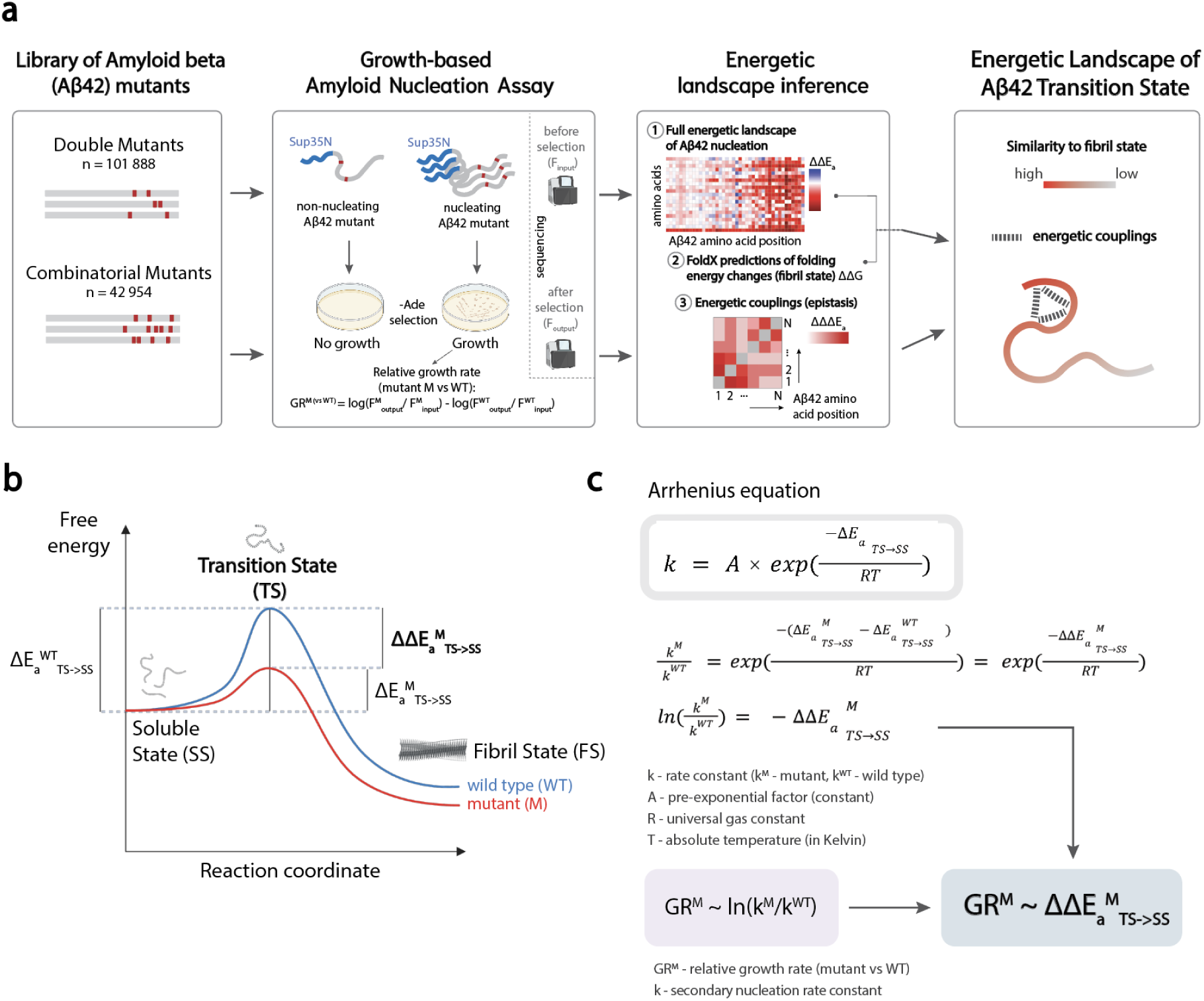
Quantifying changes in amyloid nucleation reaction activation energies at scale. **a** Schematic overview of the study: mutant libraries are selected through an assay where cell growth depends on amyloid nucleation and is quantified by sequencing. The resulting relative growth rates are used to fit a mechanistic model to infer activation energy terms and energetic couplings which can be used to map the energetic landscape of the Aβ42 transition state. **b**, Free energy landscape of the amyloid nucleation reaction. **c**, Arrhenius equation and derivation of linear relationship between measured relative growth rates and change in activation energy; k - rate constant (k_M_ for mutant, k_WT_ for wildtype), A - pre-exponential constant, ΔE_a_-activation energy, R - universal gas constant, T - absolute temperature (Kelvin), TS - transition state, FS - fibril state, SS - soluble state, ΔΔE ^M^- change in activation energy for mutant M (vs WT), GR^M^ - relative growth rate (for mutant M vs WT), к - nucleation rate constant.

One disruptive approach for probing transition states, pioneered by Fersht and colleagues, is to use mutations^18–20^. Mutations that stabilise a transition state (or increase the energy of the initial soluble state) will lower E_a_ and accelerate a reaction whereas mutations that destabilise the transition state will slow it. Kinetic measurements therefore allow the importance of individual residues and potential structural contacts in a transition state to be probed. Moreover, comparing changes in kinetics to changes in stability can provide information on the degree of native structure around a mutated residue in a transition state^17,20^.

Using this approach, the transition states of enzymes and protein folding pathways have been probed using a small number of individual mutations^21–23^, providing important structural insights^18,19^. More recently ten mutations were used to probe the transition state of an amyloid elongation reaction, revealing that monomers that successfully bind to the fibril ends have fibril-like contacts^24^. Another study combined molecular dynamic simulations with kinetic experiments to suggest a hairpin trimer structure for the transition state of Tau amyloids^25^. However, in all cases, due to the difficulty of making kinetic measurements, only a very small number of carefully chosen mutations have been characterised.

More generally, the genotype-phenotype landscapes of protein aggregation reactions are largely unexplored. To address this shortcoming, we have developed methods that allow the nucleation kinetics of thousands of sequences to be quantified in parallel using pooled mutation-selection-sequencing experiments^26–29^. This has opened up the possibility of exploring the genetic architecture of amyloid formation —the set of rules that govern how mutations combine to determine the rate of fibril assembly. It could be that amyloid genotype–phenotype landscapes are complex, requiring many parameters to accurately predict changes in nucleation when mutations are combined. Alternatively, nucleation genetics might be simple. We consider a genotype-phenotype landscape simple if it can be described using few parameters^30^ (providing a large data compression) and parameters that are interpretable (providing understanding).

Here we directly tackle the challenge of exploring amyloid genotype-phenotype landscapes, using Aβ42 as a model system. In total we quantify the nucleation rates of >140,000 combinatorial mutants of Aβ42, including >100,000 double mutants and >40,000 higher order combinations of mutations. Fitting energy models to this dataset using neural networks reveals a simple genetic architecture, with nucleation rates accurately predicted by additive changes in the reaction activation energy and a contribution from rare pairwise energetic couplings. These simple energy models represent large compressions of the full genotype space - 102-fold and 1581-fold for the models with and without energetic couplings, respectively. The strongest energetic couplings between mutations are, moreover, not random but strongly enriched between a subset of residues that form structural contacts in mature Aβ42 fibril structures. A simple interpretation of the energy landscape is that these contacts are already formed in the transition state of the nucleation reaction, predicting a structural model for the Aβ42 nucleation transition state. Using this approach it should be possible to massively perturb the energy landscapes of additional nucleation reactions, and, in combination with additional selection assays, the energetics of protein transition states more generally.

## Results

### Quantifying the nucleation of >100,000 variants of Aβ42

The rate of an amyloid nucleation reaction depends exponentially on the activation energy (E_a_ ) of the reaction, as described by the Arrhenius equation 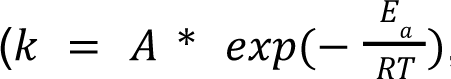, where *k* is the nucleation rate constant, *A* is a pre-exponential factor, *T* is the absolute temperature, and *R* is the universal gas constant, Fig. 1b,c)^31–33^. To quantify changes in *k* for all mutations in amyloid beta (Aβ42) we employed a kinetic selection assay in which the rate of Aβ42 nucleation controls the growth rate of yeast cells (Fig. 1a, Fig. S1). Comparison with *in vitro* nucleation rate constants from different studies shows that relative growth rates are proportional to the logarithm of nucleation rate constants^26,27^. However, the dynamic range of the assay is bounded by the maximum and minimum quantifiable relative growth rates, limiting measurement precision.

To expand the measurement range of the selection assay we employed double mutant libraries. Testing mutations in fast nucleating variants of Aβ42 provides an expanded dynamic range for quantifying decreased nucleation, whereas testing mutations in slow nucleating Aβ42 variants expands the dynamic range for quantifying increased nucleation. Using a large number of double mutants reduces the influence of specific interactions between mutations^34,35^.

In total, we quantified the relative growth rates of 101,888 double mutants of Aβ42 using three separate libraries and triplicate selection experiments (Fig. 2a-c, Fig. S2b, Supplementary Table 2 and Methods). Across these double mutants, the effect of each amino acid (AA) substitution is quantified in a median of 215 Aβ42 single mutants, providing measurements of changes in nucleation upon mutation in Aβ42 sequences with many different rates of nucleation.

**Figure 2.**
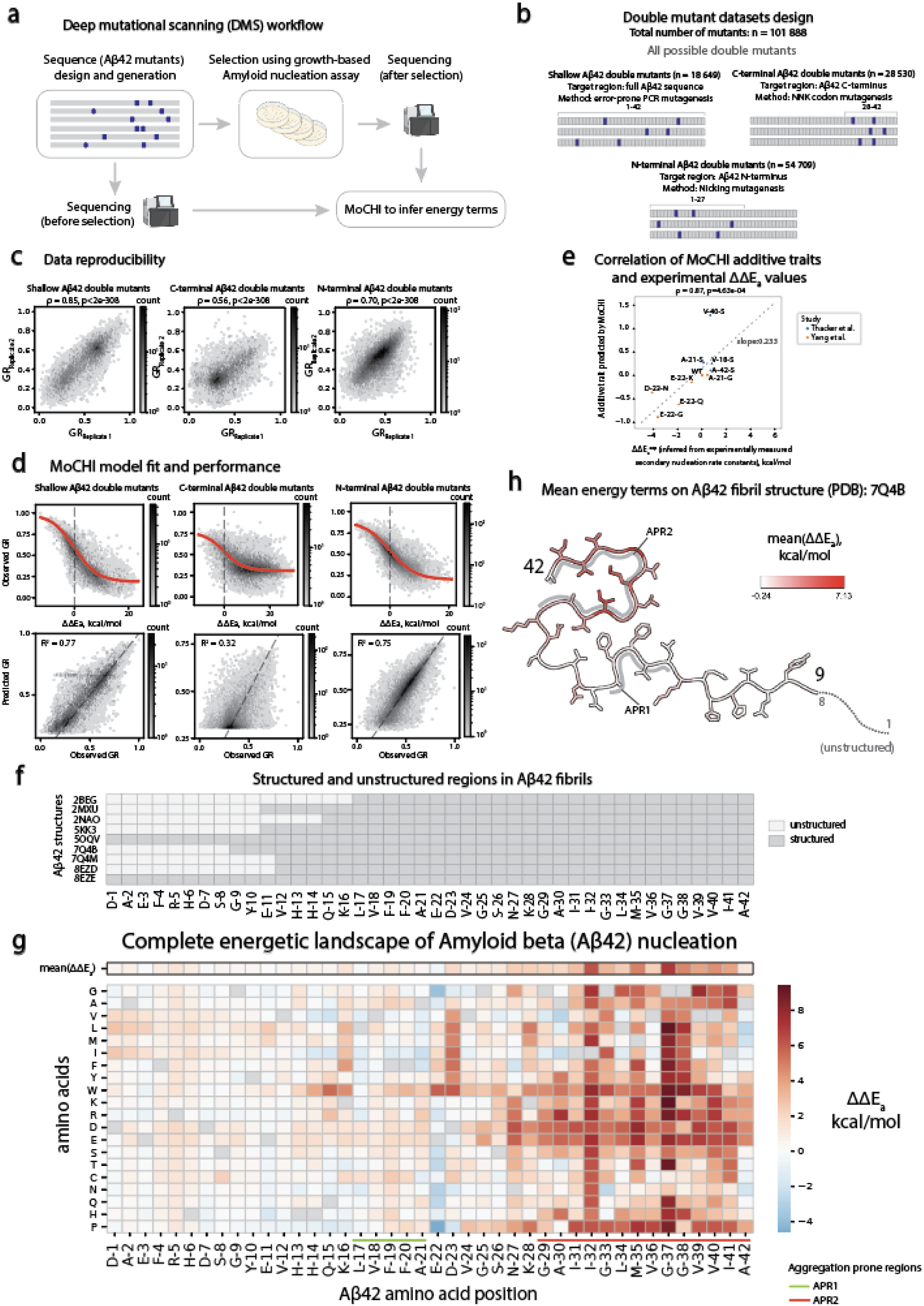
Complete activation energy landscape for Aβ42 single mutants. **a**, Schematic overview of the Deep Mutational Scanning (DMS) approach. **b**, Aβ42 double mutant libraries’ design. **c**, Inter-replicate correlations of relative growth rates (GR) for shallow, C-terminal and N-terminal Aβ42 double mutant libraries (from left to right); Spearman’s ρ (correlation) coefficients and associated p-values are reported. **d**, MoCHI model fit (top panels, red trend line represents sigmoid function fit, vertical dashed black line indicate 0 in x axis) and correlation between observed and predicted relative growth rates (bottom panels) for shallow, C-terminal and N-terminal Aβ42 double mutants datasets. **e**, Scatterplot of additive trait values predicted by MoCHI^35^ model trained on double mutant datasets (Y axis) and experimentally-derived ΔΔE_a_ values (X axis, derived from secondary nucleation rate constants) used to calibrate MoCHI-inferred terms to kcal/mol units (see Methods for details). Dashed grey line represents linear regression fit for the data. **f**, Overview of structured and unstructured regions in Aβ42 fibrils. **g**, Complete energetic landscape of Aβ42 nucleation: heatmap of inferred activation energy terms (ΔΔE_a_) for all possible substitutions in Aβ42. Mean ΔΔE_a_ values for each position are displayed in the top row of the heatmap (outlined in black). Aggregation prone regions 1 (APR1) and 2 (APR2) are highlighted in light green and orange, respectively. **h**, Cross section of 7Q4B PDB structure of Aβ42 fibrils with residues coloured by mean ΔΔE_a_ per position. Aggregation prone regions 1 (APR1) and 2 (APR2) are highlighted in grey.

### Changes in activation energy for all substitutions in Aβ42

We used MoCHI, a flexible toolkit for fitting models to deep mutational scanning data^34,35^ to infer the change in activation energy, ΔΔE_a_, for all possible AA substitutions. The model assumes additivity of free energy changes across the entire double mutant dataset, with a sigmoidal function accounting for the upper and lower bounds of the relative growth rate measurements (Fig. 2d, Fig. S2c). The resulting inferred free energies were then scaled linearly using *in vitro* measurements^36,37^ (Fig. 2e, Fig. S2a and Methods). Adding the inferred free energy changes for each mutation provides very good prediction of the double mutant nucleation rates (Pearson’s R^2^ = 0.72, evaluated by ten-fold cross validation, total of 60,510 variants) (Fig. 2d, Supplementary Table 3). The very good performance of the model highlights a non-trivial feature of the amyloid genetic landscape: the rate of nucleation of double mutants can be predicted by simple additivity of the energetic effects of single mutants (Fig. 2d).

To our knowledge, this is the first complete map of changes in activation energy for mutations in any protein (Fig. 2g). The map reveals many interesting features of the mutational landscape of Aβ42. In total, 720 AA substitutions (86%) affect the nucleation activation energy (ΔΔE_a_ ≠ 0 kcal/mol, Z-test, FDR < 0.05, Benjamini-Hochberg correction) including in all positions of Aβ42. 605 substitutions (72%) increase the activation energy (ΔΔE_a_ > 0 kcal/mol, FDR < 0.05) to slow nucleation and 115 (14%) decrease ΔΔE_a_ (ΔΔE_a_ < 0 kcal/mol, FDR < 0.05) to increase nucleation (Fig. 2g). 482 substitutions (57%) cause a larger than 1 kcal/mol change in ΔE_a_, with 443 increasing and 39 decreasing ΔE_a_ by > 1 kcal/mol. The energy changes are very well correlated with the relative growth rates of individual substitutions (ρ = 0.94, p < 2e-308, Fig. S3a, Fig. 2e), further highlighting the additive nature of the assay, but with an expanded dynamic range for mutations that slow nucleation (Fig. S3a-d, Supplementary Table 4, particularly mutations of G-37, G-38, L-34 and M-35).

### Importance of the C-terminus for Aβ42 nucleation

The complete activation energy landscape reveals a striking asymmetry in Aβ42: mutations towards the C-terminus (residues 29 to 42) have much larger effects on the activation energy than mutations before residue 29 (Fig. 2g,h, Fig. S2d). Not all residues are structurally resolved in mature Aβ42 fibrils^12,38–43^, but this distinction between structured and unstructured positions does not explain the difference in activation energies as mutations in many residues that are structured in mature fibrils have only small effects on the activation energy (Fig. 2f,g, Fig. S3e).

The primary sequence of Aβ42 contains two hydrophobic regions: residues 17–21 (previously referred to as aggregation prone region 1, APR1) and residues 29–42 (APR2)^27,44–46^. Both APR1 and APR2 form hydrophobic cores in mature fibril structures (Fig. 2f,h, Fig. S2d) and both have been proposed to be important for fibril stability^46^. In our data, however, mutations in APR2 have much stronger effects on E_a_ than mutations in APR1 (Fig. S2e), indicating that APR2 is more important for the rate-limiting step of the Aβ42 nucleation reaction.

A parsimonious hypothesis based on these comprehensive activation energy measurements therefore is that the C-terminus of Aβ42—but not more N-terminal residues that are also structured in mature fibrils —constitutes the structured region of the nucleation transition state.

### Comparison of activation energy changes to mature fibril stability changes

We next compared the importance of residues for nucleation to their importance for the stability of mature fibrils. For each mature fibril polymorph of Aβ42 we calculated the predicted effect of every mutation on the thermodynamic stability of the fibril as the change in Gibbs free energy, ΔΔG, (see Methods, Fig. 3a, Fig. S4)^24,46^. The stability energy matrices show that both APR1 and APR2 are important for structural stability across polymorphs, with mutations in both hydrophobic cores reducing thermodynamic stability (Fig. 3b, Fig. S5i).

**Figure 3.**
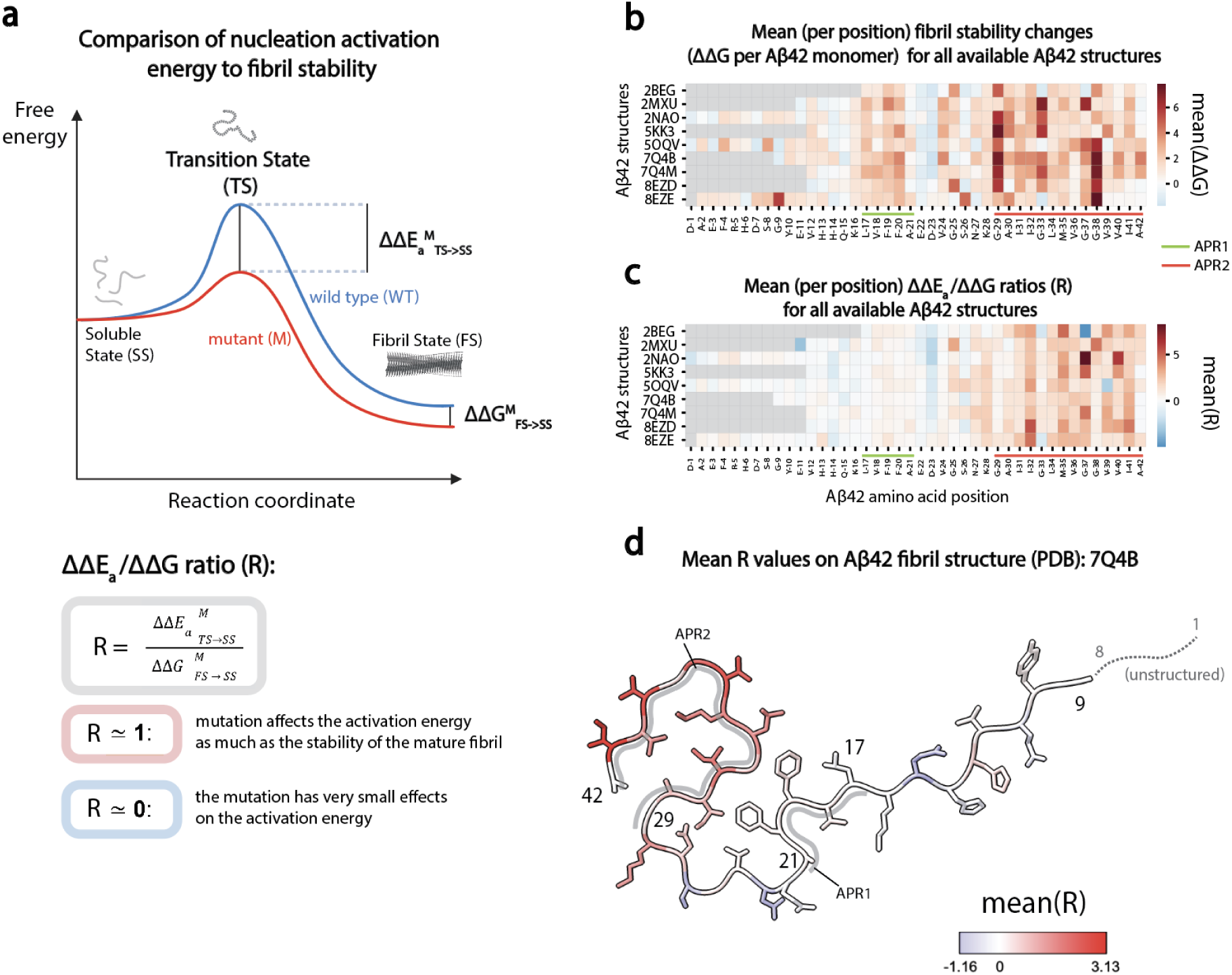
Comparing changes in activation energy to changes in mature fibril stability. **a**, Free energy landscape of the amyloid nucleation reaction (top). Changes in activation energy (ΔΔE_a_^M^_TS→SS_) and in fibril stability (ΔΔG^M^_FS→SS_) upon mutation (M) are highlighted. Principles of ΔΔE_a_/ΔΔG ratio analysis (bottom). **b**, Mean (per position) fibril stability changes (ΔΔG per Aβ42 monomer) as predicted by FoldX for all available Aβ42 structures for those substitutions that are later considered in ratio analysis (ΔΔG values are selected using the following criteria: 0.6 kcal/mol < |ddG| < 10 kcal/mol, see Methods). Aggregation prone regions 1 (APR1) and 2 (APR2) are highlighted in light green and orange, respectively. **c**, Mean (per position) ΔΔE_a_/ΔΔG ratios (R) for all available Aβ42 structures. Aggregation prone regions 1 (APR1) and 2 (APR2) are highlighted in light green and orange, respectively. **d**, Cross section of 7Q4B PDB structure of Aβ42 fibrils with residues coloured by mean R-values per position.

To more formally compare the changes in activation energy, ΔΔE_a_, to changes in fibril stability, ΔΔG, we calculated the ratio of ΔΔE_a_ to ΔΔG, an approach inspired by phi-value analysis^47–51^. A ΔΔE_a_/ΔΔG ratio of one indicates that a mutation affects the activation energy as much as the stability of the mature fibril, whereas a ratio near zero means the mutation has very small effects on the activation energy. When mutations only mildly perturb a reaction path, ΔΔE_a_/ΔΔG ratios can be interpreted as quantifying the degree of structural conservation between the transition state and the mature fibril (Fig. 3a)^47–51^.

Calculating ΔΔE_a_/ΔΔG ratios for all moderate effect mutations (see Methods) and all mature fibril structures of Aβ42 shows that the changes in activation energy are similar to changes in stability for mutations in APR2 but much smaller than changes in stability for mutations in APR1 (Fig. 3b-d, Fig. S5a-h, Fig. S6c, Supplementary Table 6, Supplementary Table 7). A simple interpretation of this result is that APR2 has a similar structure in the Aβ42 transition state as in mature fibrils, whereas APR1 does not.

The mature Aβ42 fibril structure with ΔΔE_a_/ΔΔG ratios closest to 1 (smallest root mean square distance to 1 across all ratios, Supplementary Table 8) in the APR2 region is 7Q4B, which is a cryo-EM structure of Aβ42 fibrils from familial Alzheimer’s disease brains^12^, suggesting the structured region of the Aβ42 transition state is most similar to this mature fibril polymorph. However, most Aβ42 mature fibril polymorphs are structured very similarly in the C-terminal APR2 region (Fig. S5a-h), and the ΔΔE_a_/ΔΔG ratios are consistently high in this region across the polymorphs (Fig. 3c, Fig. S6a,b).

In summary, a parsimonious interpretation of the comprehensive activation energy measurements and comparisons to the stability of mature fibrils is that APR1 is largely unstructured in the Aβ42 nucleation transition state, whereas APR2 is structured and in a manner that is similar to the structure of this region in mature fibrils (Fig. 3c,d).

### Massively parallel combinatorial double mutant cycles

To further probe the energy landscape of the Aβ42 nucleation reaction, we performed double mutant cycles^18,19^. In double mutant cycles, the energy changes in double mutants are compared to those predicted by summing the energy changes in single mutants. For thermodynamic stability, the resulting energetic couplings can provide structural information : while mutations in non-contacting positions normally result in additive changes in free energy, mutations in structurally contacting residues are often energetically coupled, causing non-additive changes in energy when combined. Quantifying energy changes in double mutants and comparing these to energy changes in single mutants therefore provides a powerful approach to probe the structures of proteins, and this approach can also be applied to probe the structures of short-lived high energy states such as transition states^18,19,52,53^.

We quantified energetic couplings between mutations in Aβ42 by designing combinatorial mutagenesis libraries in which the effects of single and double mutants were quantified in many different variants of Aβ42 with different nucleation rates (Fig. 4a,b,d, Fig. S7a). In total we quantified the nucleation of 42,954 combinatorial variants containing up to 8 different mutations (Fig. 4c). Each specific pair of mutations is therefore tested in a median of 1,024 different genotypes of Aβ42, allowing accurate measurements of both free energy changes and the energetic interactions between mutations (the energetic couplings). As for our comprehensive single mutant energy measurements, quantifying double mutants in Aβ42 variants with fast and slow nucleation rates allows more precise inference of changes in activation energy and measurement across an expanded dynamic range. To reduce the chances of inducing large structural changes along the reaction path we deliberately employed mutations that conserve side chain hydrophobicity (Fig. 4a and Methods). The growth rates of the combinatorial mutants were well correlated between replicate experiments (Fig. 4b). The genotype frequency landscape also matched expectations, with most genotypes in the library containing 6 substitutions from the wild-type Aβ42 (Fig. 4c). The median growth rates of multi-mutants decreased with an increasing number of mutations, but combinatorial mutants of all orders still had a wide range of rates (Fig. 4c).

**Figure 4.**
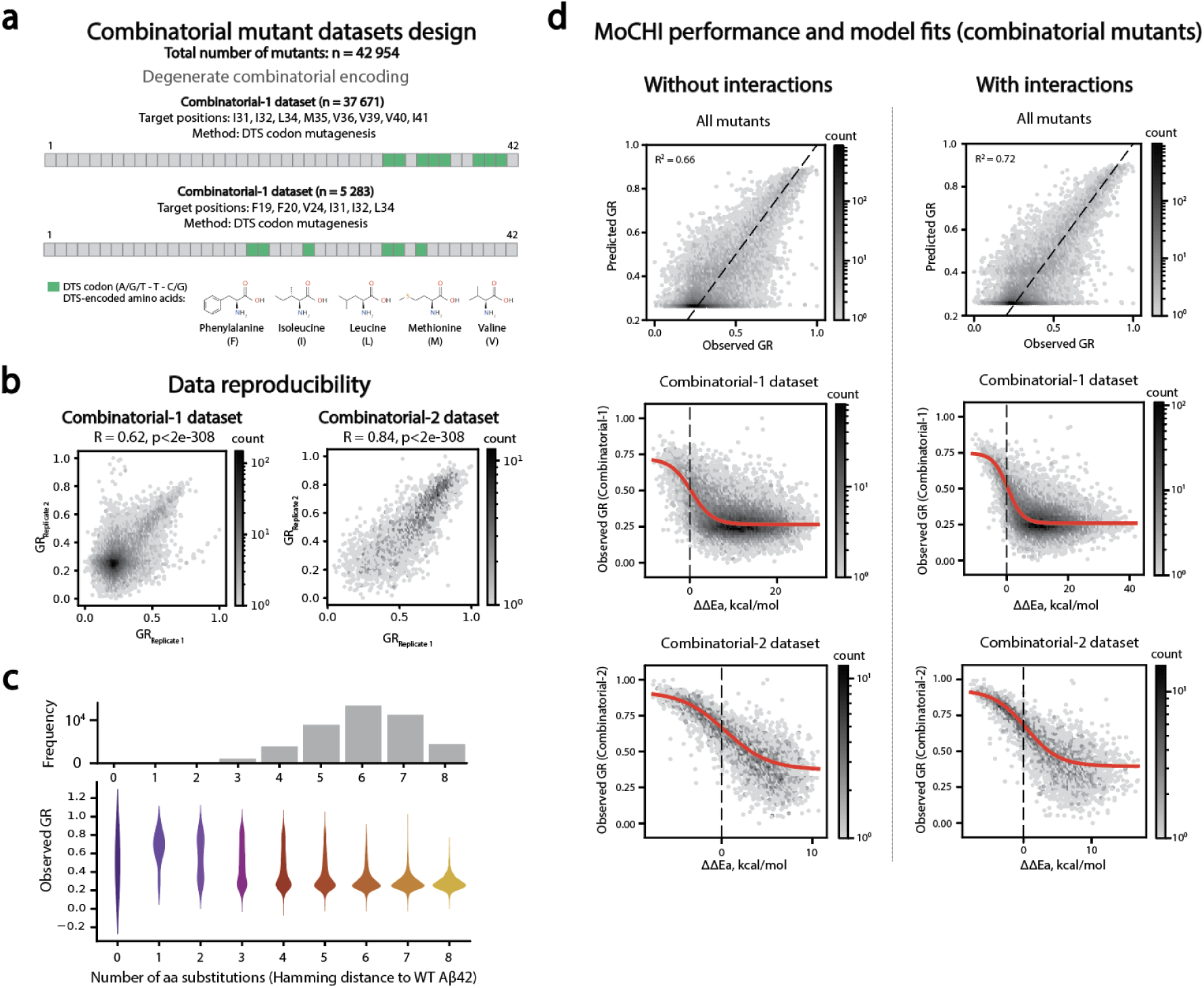
Simple genetic and energetic architecture of Aβ42. **a**, Combinatorial Aβ42 mutant libraries’ design; DTS codons encode Phenylalanine (F), Methionine (M), L (Leucine), I (Isoleucine), V (Valine). **b**, Inter-replicate correlations of relative growth rates (GR) for Combinatorial-1 (left) and Combinatorial-2 (right) Aβ42 combinatorial mutant libraries. Pearson’s correlation coefficients (R) and associated p-values are indicated. **c**, (top) Histogram of the number of observed combinatorial mutants of Aβ42 at increasing Hamming distances from the wild-type Aβ42 (denoted by WT), for which the x axis is shared with the bottom panel; (bottom) Violin plot showing distributions of relative growth rates of combinatorial mutants of Aβ42 inferred from deep sequencing data versus number of aa substitutions. **d**, Left column - MoCHI performance (top row, correlation between observed and predicted relative growth rates) for all combinatorial mutants and fits (middle and bottom row, red trend line represents sigmoid function fit, vertical dashed black line indicates 0 in x axis) for Combinatorial-1 and Combinatorial-2 Aβ42 combinatorial mutants for model that does not account for interactions (energetic couplings); Right column: MoCHI performance (top row, correlation between observed and predicted relative growth rates) for all combinatorial mutants and fits (middle and bottom row, red trend line represents sigmoid function fit, vertical dashed black line indicates 0 in x axis) for Combinatorial-1 and Combinatorial-2 Aβ42 combinatorial mutants for model that accounts for interactions (energetic couplings).

### The genetic architecture of amyloid nucleation

We used MoCHI to infer the changes in free energy (ΔΔE_a_) in multi-mutants and the energetic couplings between mutations (ΔΔΔE_a_, Fig. 4d, Fig. S7b,c). We first considered a model in which only the effects of individual mutations are measured and these combine additively in combinatorial mutants. Despite only containing one energy change per mutation (a total of 44 parameters) this model provides very good prediction of the nucleation rates for the complete dataset (Pearson’s R^2^ = 0.66, evaluated by ten-fold cross validation, total of 42,954 genotypes) (Fig. 4d, top panels). The model is a very large compression of the full genotype space (∼1581-fold, with 4^8^ + 4^6^ - 4^3^ possible genotypes vs. 44 parameters).

We next considered a model in which pairwise energetic couplings were also inferred between all pairs of genotypes. This model provided improved predictive performance, accounting for an additional ∼6% of variance (R^2^ = 0.72 vs. R^2^ = 0.66 for the model without energetic couplings). The model still represents a large data compression of ∼102-fold (4^8^ + 4^6^ - 4^3^ possible genotypes vs. 684 model parameters).

### Sparse pairwise energetic interactions

In total, the model with pairwise energetic interactions quantifies 640 energetic couplings between 40 pairs of residues (Fig. S8a-j, Supplementary Table 9). Strikingly, the vast majority of the energetic couplings are very small (median ΔΔΔE_a_ = 0.3 kcal/mol), with 535/640 (84%) having an absolute value < 1 kcal/mol (Fig. S8k).

Less than 4% of energetic couplings have an absolute value > 2 kcal/mol, with 15 positive and 9 negative couplings. This is consistent with the expectation of energetic additivity for most combinations of mutations and the small improvement in predictive performance of the model over a completely additive model (Fig. 4d).

### Energetic couplings predict a model for the Aβ42 nucleation transition state

We used the average of the absolute couplings ΔΔΔE_a_ between different mutations in the same residues as an overall metric of the energetic non-independence of two sites (interaction score)^54^. The median of the interaction score distribution is 0.53 kcal/mol, with only 4/40 pairs of residues having an average coupling > 1 kcal/mol (Fig. 5a, Fig. S8l). Very interestingly, all four of these strongly coupled sites are located between residues in the APR2 region: M35-V36, I32-M35, I32-V40, M35-V40.

**Figure 5.**
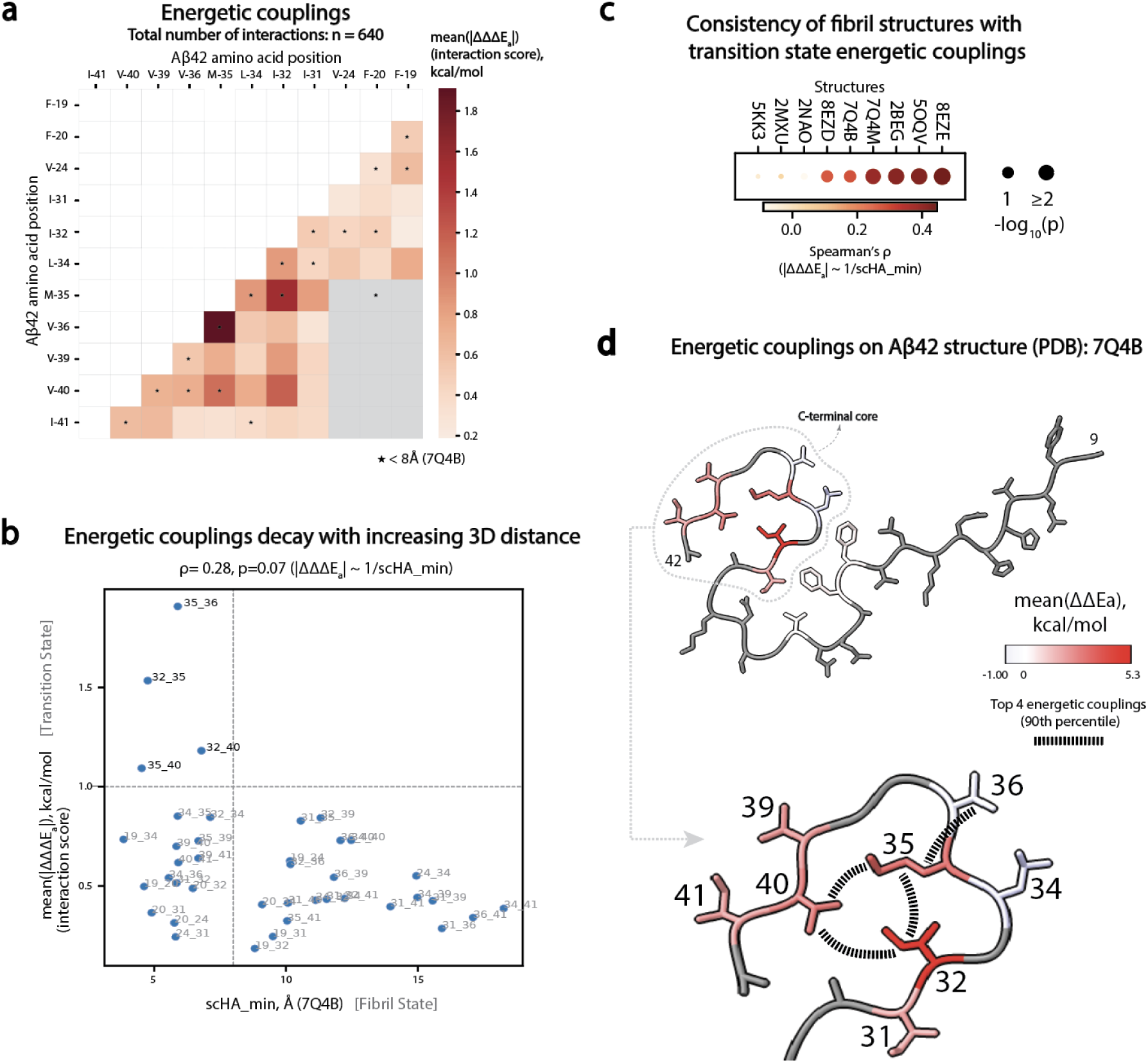
Energetic interactions in Aβ42 nucleation. **a**, Interaction scores (mean absolute value of energetic couplings for a pair of positions, mean(|ΔΔΔE_a_|)) for mutagenised Aβ42 position pairs. Star symbols (⭑) indicate residue pairs closer than 8 Å in 7Q4B Aβ42 structure. **b**, Scatterplot of interaction scores for pairs of positions and inter-residue distance for the corresponding pairs of amino acids in 3D space (scHA_min, minimum heavy atom side chain distance) as measured in the Aβ42 fibril structure 7Q4B; dashed light grey vertical line marks 8 Å, dashed light grey horizontal line marks interaction score (mean(|ΔΔΔE_a_|) of 1 kcal/mol. **c**, Dotplot representing consistency of transition state interactions with fibril structures (2BEG, 2MXU, 2NAO, 5KK3, 5OQV, 7Q4B, 7Q4M, 8EZD, 8EZE) according to energetic couplings and amino acid inter-residue distances; size of the dot reflects statistical significance (-log_10_(p)) of the correlation between interaction score (mean(|ΔΔΔE_a_|)) and inverse of the inter-residue distance (1/scHA_min); colour of the dot represents Spearman’s ρ (correlation) coefficient. **d**, Cross section of 7Q4B PDB structure of Aβ42 fibrils with residues coloured by mean activation energy terms (ΔΔE_a_) inferred from the MoCHI model trained on combinatorial mutants datasets (residues with no inferred ΔΔE_a_ are shown in grey). Positions mutagenised in combinatorial datasets (Combinatorial-1 and Combinatorial-2) are labelled on the PDB structure. Top 4 interacting position pairs (in 90^th^ percentile of interaction score distribution) are connected with dashed black lines.

To compare the energetic couplings to the structures of mature fibrils, we plotted the mean absolute ΔΔΔE_a_ values against the distance between amino acid residues in 3D space (minimal side chain heavy atom distance, scHA_min) separately for each of nine Aβ42 fibril polymorphs (Fig. 5b, Fig. S9a, Supplementary Table 10). The couplings were correlated with the inverse of the distances in four polymorphs (8EZE, 5OQV, 2BEG, 7Q4M, p < 0.02) (Fig. 5c). In Fig. 5b we show the plot of interaction scores vs. 3D structural distance for fibrils extracted from familial Alzheimer’s disease brains (PDB 7Q4B)^12^ which has a striking L-shaped distribution of coupling strength versus 3D distances, with a subset of structural contacts in the mature fibrils strongly energetically coupled. Visualising these energetic couplings on the mature fibril structure further highlights that a subset of spatially-clustered structural contacts in the C-terminal APR2 region of mature fibrils are strongly energetically coupled when they are mutated (Fig. 5d, Fig. S9b).

Changes in the activation energy of a reaction can be caused by changes in the energy of the transition state, changes in the energy of the starting state, or both (Fig. 1b). For the Aβ42 nucleation reaction, therefore, the energetic effects of mutations and the energetic interactions between them could be driven by changes in the energy of both the transition state or the starting soluble state. However, that all of the most strongly interacting pairs of positions match structural contacts in the C-terminal APR2 region of mature fibril structures suggests a parsimonious explanation of our data: mutations and their interactions primarily affect the energy of the transition state and this transition state is already partially structured in the C-terminal APR2 region of Aβ42. Applying Occam’s razor therefore allows us to propose a structural model for the Aβ42 nucleation transition state in which only the C-terminal APR2 region is structured and similarly to the structure of mature amyloid fibrils.

## Discussion

We have demonstrated here the feasibility of using DNA synthesis-selection-sequencing experiments to massively perturb the energy landscape of an amyloid nucleation reaction and we have used the resulting data to propose a parsimonious model for the transition state of a reaction that initiates Aβ42 aggregation and so, potentially, Alzheimer’s disease.

In total, we measured the nucleation rates of >140,000 protein sequences. Using neural networks to fit energy models allowed us to quantify the change in nucleation activation energy for all 798 amino acid substitutions in Aβ42 and, in addition, the energetic interactions between 640 pairs of mutations. This is, to our knowledge, the first comprehensive measurement of changes in activation energy for any protein and also the first large-scale measurement of energetic couplings for a kinetic process. Our work builds on pioneering studies by Fersht and others^17,23,24^, expanding activation energy and energetic coupling measurements to the scale of all mutations in a sequence. The resulting energy models are simple and interpretable and represent large compressions of the genotype landscape. However they also provide excellent predictive performance of the nucleation rates of the entire experimental dataset, revealing the reassuringly simple genetic architecture of an amyloid nucleation reaction.

We believe that the strategy taken here can now be applied to massively perturb the energy landscapes of additional amyloid nucleation reactions^29^, as well as thousands of short nucleating peptides^28^. A key feature of our approach is the use of a kinetic selection assay where enrichments report on the rate of a reaction. This is not true for most synthesis-selection-sequencing experiments, where enrichments typically depend on thermodynamic stabilities^34,55,56^. An important challenge for the future is the development of pooled kinetic selections for reactions other than amyloid nucleation. Approaches using microfluidics^57^ and droplet-based selections^58,59^ may facilitate this, allowing massive perturbation of the energy landscapes of additional protein reactions and high-resolution analysis of the structures of protein transition states.

Another key component of our approach is the use of double mutants and combinatorial mutagenesis. Quantifying mutational effects and interactions in variants with faster and slower kinetics expands the dynamic range of an assay and so the precision of energy measurements. The approach assumes that changes in free energy are largely additive when mutations are combined. The good predictive performance of our models even when many different mutations are combined, and the consistency of non-additive energetic couplings with structural contacts in a region of mature fibril atomic structures suggests that this is indeed the case for Aβ42 nucleation.

Our comprehensive energy landscape identifies the residues which, when mutated, have the largest effect on the activation energy of the Aβ42 nucleation reaction and also quantifies the energetic coupling between them. Changes in the activation energy of a reaction can be caused by changes in the energy of the transition state or the starting state (Fig. 1b). Formally we cannot distinguish between these two possibilities as explanations for our data. However, we believe that the consistency between the energetic interaction map and the structure of a region of mature fibrils (Fig. 3) strongly suggests that mutations are primarily altering the energy of the transition state and that this transition state is already partially structured in the APR2 region of Aβ42 (Fig. 2).

Amyloid fibrils of Aβ42 are a defining pathological hallmark of all forms of Alzheimer’s disease and variants in Aβ42 also cause familial Alzheimer’s disease^3^. Moreover, the only therapeutics demonstrated to slow the progression of Alzheimer’s disease in clinical trials are antibodies targeting Aβ42^60,61^. Understanding the reaction that initiates the formation and spread of Aβ42 fibrils is a central goal of Alzheimer’s research: this reaction is the key process to stop in order to prevent the disease. We believe our data and our data-driven model of the Aβ42 transition state is an important step forward in this endeavour.

## Methods

### Library designs and construction

#### N-terminal Aβ42 double mutant library

To obtain the N-terminal Aβ42 double mutant library (532 possible single aa mutants and 151,200 possible double aa mutants) we used a nicking mutagenesis approach described in Wrenbeck et al.^62^ . The Aβ42 region with 25 bp upstream and 21 bp downstream was amplified from plasmid PCUP1-Sup35N-Aβ42^63^ kindly provided by the Chernoff lab using oligos AB_TS_003-004 (Supplementary Table 11) to introduce AvrII and HindIII restriction sites. The target region for mutagenesis was then digested and ligated (T4 DNA Ligase, Thermo Scientific) into the nicking plasmid pGJJ057, previously digested with the same restriction sites.

There are 4 main steps in the nicking mutagenesis protocol: i) obtention of ssDNA template, ii) synthesis of a mutant strand by annealing and extension of mutagenic oligos (oligos AB_TS_005-032, Supplementary Table 11), iii) degradation of the wildtype (WT) template strand and iv) synthesis of the 2nd mutant strand (oligo AB_TS_033, Supplementary Table 11). We used 28 mutagenic oligos synthesised by IDT, each of them containing one NNK (N=A/T/C/G; K=T/G) degenerate codon at every targeted position for mutagenesis, flanked by 7 upstream and 7 downstream wild-type codons.

In order to obtain a double mutant library, we ran the protocol twice. In the first round, we obtained a library with 66% of single amino acid mutant and 33% of WT sequences (estimated by sanger sequencing of individual clones). In the second round - in which we used the single mutant library as template - we obtained a library with 66% of double and 33% of single amino acid mutant sequences.

As described in Wrenbeck et al.^62^, the N-terminal Aβ42 double mutant library was finally transformed into 10-beta Electrocompetent *E. coli* (NEB). Cells were recovered in SOC and plated on LB with ampicillin to assess transformation efficiency. A total of 2.88 million transformants were estimated, representing each variant of the library more than 18x. 50 ml of overnight culture were harvested to purify the library in the nicking plasmid pGJJ057 (QIArep Miniprep Kit, Qiagen).

In order to clone the library back inside the PCUP1-Sup35N plasmid for selection, the library in the nicking plasmid was digested with EcorI and XbaI restriction enzymes (Thermo Scientific) for 4 hours at 37°C and purified from a 2% agarose gel (QIAquick Gel Extraction Kit, Qiagen). At this stage, the purified product was ligated into the previously digested PCUP1-Sup35N and transformed into 10-beta Electrocompetent *E. coli* (NEB). For this library, a total of 3.1 million transformants were estimated.

#### C-terminal Aβ42 double mutant library

To obtain a library of double mutants in the Aβ28-42 region (285 possible single aa mutants and 37,905 possible double aa mutants), we used a NNK codon mutagenesis approach. We ordered an oligo pool (IDT) containing 105 oligos of Aβ28-42 (15 AA positions), each with 2 positions containing an NNK (N=A/T/C/G; K=T/G) degenerate codon (oligo pool AB_TS_034, Supplementary Table 11). Each NNK degenerate codon encoded 32 possible codons, resulting in a library of 107,520 unique nt sequences. Each oligo also contained 5’ (CTTTGCAGAAGATGTGGGTTCAAAC) and 3’ (TAATCTAGAGCGGCCGCCACC) constant regions for cloning purposes.

500 ng of the oligo pool were amplified by PCR (Q5 high-fidelity DNA polymerase, NEB) for 10 cycles with primers annealing to the constant regions (oligos AB_TS_037,039, Supplementary Table 11). The PCR product was incubated with ExoSAP-IT (Thermo Fisher Scientific) at 37C for 1h and purified by column purification (MinElute PCR Purification Kit, Qiagen). This library did not contain the EcoRI restriction site between Sup35N and Aβ42, which was previously used for cloning of the shallow and N-terminal Aβ42 double mutant libraries.

In parallel, empty plasmids PCUP1-Sup35N-Aβ(1-27) (i.e. missing a specific Aβ region for mutagenesis) was constructed by PCR linearisation PCUP1-Sup35N-Aβ42 using oligos AB_TS_040-041 (Supplementary Table 11). These was then linearised by PCR for 35 cycles (oligos AB_TS_044-045, Supplementary Table 11), treated with DpnI (FastDigest, Thermo Scientific) overnight and purified from a 1% agarose gel (QIAquick Gel Extraction Kit, Qiagen).

The purified library was then ligated into 200 ng of PCUP1-Sup35N-Aβ(1-27) in a 10:1 (library:plasmid) ratio by Gibson assembly with 3h of incubation. The resulting product was then dialysed for 3h by using a membrane filter (MF-Millipore 0.025 µm membrane, Merck) and concentrated 10X using a speed vacuum concentrator. Finally, the library was transformed into 10-beta Electrocompetent *E. coli* (NEB). Cells were recovered in SOC and plated on LB with ampicillin to assess transformation efficiency. 50 ml of overnight culture were harvested to purify the plasmid library (QIArep Miniprep Kit, Qiagen). A total of 1.8 million transformants were estimated, representing each variant of the library more than 47 times.

#### Shallow Aβ42 double mutant library

We built a library containing double mutants across the entire Aβ42 sequence (798 possible single aa mutants and 310,821 possible double aa mutants) in a previous study^26^. Briefly, the library was obtained by error-prone PCR and cloned by digestion and ligation into the PCUP1-Sup35N plasmid (oligos AB_TS_001-002). Additional details on library construction are provided in Seuma et al.^26^. For this library, a total of 4.1 million transformants were estimated, representing each variant in the library >10x.

#### Combinatorial Aβ42 libraries

We designed 2 combinatorial libraries (Combinatorial-1 and Combinatorial-2) by mutagenizing specific positions of Aβ1-42 (I31, I32, L34, M35, V36, V39, V40 and I41 in Combinatorial-1; F19, F20, V24, I31, I32 and L34 in Combinatorial-2) to a specific subset of hydrophobic amino acids. We used the DTS degenerate codon (D=A/T/G; T=C/G) to encode methionine, valine, leucine, isoleucine and phenylalanine. For each library, we purchased a 4 nmole ultramer from IDT (oligos AB_TS_035-036, Supplementary Table 11) covering the target region for mutagenesis (Aβ29-42 for Combinatorial-1 and Aβ12-42 for Combinatorial-2) containing DTS codons at the above mentioned positions. The oligos also contain 5’ and 3’ constant regions for cloning purposes. The number of possible unique amino acid sequences is 390,625 for Combinatorial-1 and 15,625 for Combinatorial-2.

These libraries were amplified and cloned following the same steps as for the C-terminal Aβ42 double mutant library (oligos AB_TS_037-045). The purified Combinatorial-1 library was ligated into PCUP1-Sup35N-Aβ(1-27) while Combinatorial-2 into PCUP1-Sup35N-Aβ(1-11). Combinatorial-1 and Combinatorial-2 also did not contain the EcoRI restriction site between Sup35N and Aβ42, which was previously used for cloning of the shallow and N-terminal Aβ42 double mutant libraries.

A total of 0.38 million transformants were estimated for Combinatorial-2 representing each variant of the library more than 24 times. The Combinatorial-1 library was bottlenecked to 0.94 million transformants due to high complexity.

### Yeast transformation

*Saccharomyces cerevisiae* GT409 [psi-pin-] (MATa ade1-14 his3 leu2-3,112 lys2 trp1 ura3-52) strain (kindly provided by the Chernoff lab) was used in all experiments in this study. Lithium acetate transformation was used to transform each plasmid library in yeast cells, in three biological replicates. An individual colony for each transformation tube was grown overnight in 20 ml YPDA at 30°C and 200 rpm. Once at saturation, cells were diluted to OD600= 0.25 in 175 ml YPDA and grown until exponential phase, for ∼5h. Cells were then harvested, washed with milliQ and resuspended in 8.5 ml sorbitol mixture (100 mM LiOAc, 10 mM Tris pH 8, 1 mM EDTA, 1M sorbitol). 5ug of plasmid library, 175 μl of ssDNA (10 mg/ml, UltraPure, Thermo Scientific) and 35 ml of PEG mixture (100 mM LiOAc, 10 mM Tris pH 8, 1 mM EDTA pH 8, 40% PEG3350) were added to the cells, and incubated for 30 min at RT. Heat-shock was performed for 15 min at 42°C in a water bath. Cells were then harvested, washed and resuspended in 50 ml recovery medium (YPDA, 0.5M sorbitol) for 1h at 30°C 200 rpm. Finally, cells were harvested, washed and resuspended in 350 ml plasmid selection medium (-URA 10% glucose) and grown for 48h. Once at saturation, cells were diluted in 200 ml plasmid selection medium (-URA 10% glucose) to OD600= 0.05 and grown until they reached the exponential phase, for ∼15h. Lastly, cells were harvested and stored at -80°C in 25% glycerol. A minimum of 3.7, 3, 2.2, 0.99 and 0.43 million transformants were estimated for each of the three biological replicates of shallow Aβ42 double mutant, N-terminal Aβ42 double mutant, C-terminal Aβ42 double mutant, Combinatorial-1 and Combinatorial-2 combinatorial libraries, respectively.

### Selection experiments

Each transformation replicate was later used for selection. Tubes were thawed from -80C, washed and resuspended in 100-1000ml plasmid selection medium (-URA 10% glucose) - ensuring 100 cells each variant in the library - at OD=0.05 and grown until exponential at 30°C 200 rpm. Once at exponential, cells were diluted in 100-1000 ml protein induction medium (-URA 10% glucose, 100μM Cu2SO4) at OD=0.05 and grown for 24 h at 30°C 200 rpm. After 24h, 25-50 ml of cells were harvested for input pellets and a minimum of 18.5, 75, 12, 230 and 1.5 million cells / replica (for shallow Aβ42 double mutant, N-terminal Aβ42 double mutant, C-terminal Aβ42 double mutant, Combinatorial-1 and Combinatorial-2 libraries, respectively) were plated on -URA-ADE selection medium plates (145cm2, Nunc, Thermo Scientific) and incubated for 6 days (for C-terminal Aβ42 double mutant, Combinatorial-1 and Combinatorial-2 combinatorial libraries) or 7 days (shallow and N-terminal Aβ42 double mutant libraries) at 30°C. The number of cells plated ensures a minimum coverage of 10 times each variant in the library. Finally, colonies were scraped off the plates and harvested for output pellets. Inputs and output pellets were stored at -20°C and later treated for DNA extraction.

### DNA extraction and sequencing library preparation

Input and output pellets (3 biological replicates each, 6 tubes in total) were resuspended in 2 ml extraction buffer (2% Triton-X, 1% SDS, 100 mM NaCl, 10 mM Tris pH 8, 1 mM EDTA pH 8) and subjected to two cycles of freezing-thawing in an ethanol-dry ice bath and water bath at 62°C for 10 min each. 1.5 ml of phenol:chloroform:isoamyl 25:24:1 was then added to the pellets and vortexed for 10 min. The aqueous phase was recovered by means of 30 min centrifugation at 3000 rpm. DNA was precipitated with 1:10 V NaOAc 3M and 2.2 V of 99% cold ethanol and incubated at -20C for 2h. After centrifugation, pellets were dried overnight at RT. The next day, pellets were resuspended in TE 1X buffer (10mM Tris pH8, 1mM EDTA pH8) and treated with 10 μl RNase A (Thermo Scientific) for 1h at 37°C. DNA was purified using a silica beads extraction kit (QIAEX II Gel Extraction Kit, Qiagen). Plasmid library concentration was quantified by quantitative PCR using primers annealing to the origin of the replication site of the plasmid (oligos AB_TS_046-047, Supplementary Table 11).

### Sequencing library preparation

Each sequencing library was prepared in a two-step PCR (Q5 high-fidelity DNA polymerase, NEB). First, the mutagenised Aβ region was amplified for 15 cycles with frame-shifted primers (oligos AB_TS_048-080, Supplementary Table 11), using 300M (for Shallow Aβ42 double mutant library), 100M (for N-terminal Aβ42 double mutant, C-terminal Aβ42 double mutant and Combinatorial-1 combinatorial libraries) and 50M (for Combinatorial-2 combinatorial library) molecules as template for each sample (3 inputs, 3 outputs). PCR products were treated with ExoSAP-IT (Thermo Fisher Scientific) at 37°C for 1h, purified by column purification (MinElute PCR Purification Kit, Qiagen) and eluted in 5 μl TE 1X buffer (10mM Tris, 1mM EDTA). 4 μl of purified product were used as template for the second PCR. In this case, samples were amplified for 10 cycles with primers containing Illumina sequencing indexes (oligos AB_TS_081-111, Supplementary Table 11). The three input samples were pooled together equimolarly, and the same for the three output samples. The final pools were purified from a 2% agarose gel using a silica beads extraction kit (QIAEX II Gel Extraction Kit, Qiagen).

Libraries were sequenced on an Illumina HiSeq2500 sequencer using 125 paired-end reads (shallow Aβ42, N-terminal and C-terminal Aβ42 double mutant libraries) or 150 paired-end sequencing on an Illumina NextSeq500 (Combinatorial-1 and Combinatorial-2 combinatorial libraries) at the CRG Genomics core facility.

### Data processing, relative growth rates and error estimates

FastQ files from paired end sequencing were processed using DiMSum^64^, an R pipeline for the analysis of deep mutational scanning data. A total of 251.6M (for Shallow Aβ42 double mutant library), 352.9M (for N-terminal Aβ42 double mutant library), 106.3M (for C-terminal Aβ42 double mutant library), 109.8M (for Combinatorial-1) and 26.8M (for Combinatorial-2, 15.9M for input samples and 10.9M for output samples) paired-end reads were obtained from sequencing.

5’ and 3’ constant regions were trimmed, allowing an error rate of 20% mismatches compared to the reference sequence. Reads were aligned and sequences with a Phred base quality score below 30 and non-designed sequences were discarded. Variants with fewer than 10 input reads in any of the replicates were discarded. Estimates from DiMSum were used to choose the filtering thresholds.

Relative growth rates and their associated error estimates (available in Supplementary Table 2) were also calculated using DiMSum. Relative growth rate for a specific variant *i*(*GR_i(vs WT)_*) in the library in each biological replicate is defined in the following way: 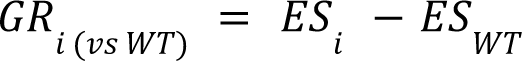, where *ES_i_* is the enrichment score for the variant *i*, and *ES_WT_* is the enrichment score of the WT Aβ42, both defined as follows: 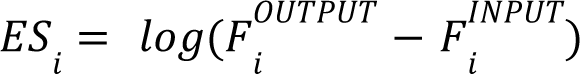

Relative growth rates for each variant were merged using error-weighted mean of each variant across replicates and centered using the error-weighted mean frequency of WT Aβ42 synonymous substitutions. Due to large possible sequence space (16-391K variants at the AA level) in combinatorial mutant datasets, WT Aβ42 was not present there, so the normalisation was done using a random variant present in those datasets (namely, GGTGCAATCATCGGATTGATGGTGGGCGGTGTGGTGTTCGCG and GTTCATCATCAAAAATTGGTGTTGTTCGCAGAAGATTTGGGTTCAAACAAAGGTGCAGTCTTGGGATTGATGGTGGGCGGTGTTGTCATAGCG were used as the WT sequence parameters in DiMSum^64^ for Combinatorial-1 and Combinatorial-2 datasets, respectively). For visualisation purposes in main figures 2,4 and supplementary figures 2, 3 and 7 relative growth rates were scaled between 0 and 1.

### Inferring changes in activation energy with MoCHI

To infer activation energy terms from relative growth rates of cells carrying nucleating Aβ42 variants, we fitted an energy model to our datasets using MoCHI^34,35^. Briefly, MoCHI takes as input amino acid sequences, measured relative growth rates and error estimates of each variant in the library. MoCHI then predicts relative growth rates for the given variants based on a particular fit (sigmoid in this case), while correcting for global non-linearities (non-specific epistasis) that in this case are due to the upper and lower limits of the growth assay. The effects of individual mutations and mutation combinations (genetic interactions) are modelled additively at the energetic level. Using the coefficients derived by the model, one then obtains the change in activation energy associated with each mutation for the phenotype of interest (in our case, amyloid nucleation).

For double mutant datasets (shallow Aβ42 double mutant library, N-terminal Aβ42 double mutant library and C-terminal Aβ42 double mutant library) we used MoCHI with default parameters for a two-state model with one phenotype (nucleation) for the three double mutants datasets, using L1 and L2 regularization with a lambda of 10^-5^ and allowing only first order (additive) energy terms. We evaluated the model using the held-out “fold” from the 10 times that the model was run on the dataset.

In order to translate (or calibrate) additive traits inferred with MoCHI^35^ into units of energy (kcal/mol), we used experimentally measured nucleation rate constants from previously published studies^37,65^ (Supplementary Table 1). We derived changes in activation energy for Aβ42 variants from the available kinetic data, using the following relationship (derived from the Arrhenius equation, see Fig. 1c): 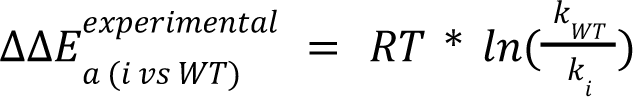, for Aβ42 variant *i*, where *R* is the universal gas constant and *T* is temperature in Kelvin (303 K, as used in MoCHI model training). For the Yang et al. dataset^37^ we used primary and secondary nucleation rate constants (k_n_ and k_2_) reported in the study to directly calculate changes in activation energy for reported Aβ42 variants (D23N, E22G, E22Q, E22K and A21G) relative to WT. For the Thacker^36^ dataset, we derived multiplicative terms k_+_k_n_ and k_+_k_2_ (primary and secondary nucleation rate constants multiplied by the rate of elongation) from the reported λ and κ values (the rate at which new fibril mass is formed via primary and secondary nucleation, respectively) and the exact model description authors provided in the supplementary material of the publication, for the reported Aβ42 variants (V18S_A21S, V40S_A42S, A-21-S, A-42-S, V-18-S and V-40-S) relative to WT, and used these multiplicative terms in the same equation (above) to calculate the activation energy change. We fitted a linear regression model to be able to predict our MoCHI-inferred additive trait values from the experimentally-derived ΔΔE_a_ values for Aβ42 variants common across the two datasets (Fig. 2e, Fig. S2a), and used the resulting slope (0.233, fitting with secondary nucleation rate-derived ΔΔE_a_ values) to calibrate the MoCHI terms to kcal/mol units.

For combinatorial mutant datasets (Combinatorial-1 and Combinatorial-2), as WT Aβ42 variant was not present in any of the two datasets, we introduced it artificially and added to the data prior to training the MoCHI model, declaring its relative growth rate as 0 and its error estimate as an arbitrary big number, 100 in this case. We used MoCHI with default parameters for a two-state model with one phenotype (nucleation) for the two combinatorial mutants datasets, using L1 and L2 regularization with a lambda of 10^-5^, and allowing first order (additive) and second order (non-additive) energy terms to account for energetic couplings. We evaluated the model using the held-out “fold” from the 10 times that the model was run on the dataset. Since we used an artificially introduced WT Aβ42 variant in MoCHI model training, we then re-centered the resulting predicted activation energy terms. To do this, we chose all common variants between double and combinatorial mutant datasets and fit a linear regression model that inferred a slope and intercept for the activation energy terms in double vs combinatorial mutant datasets. We used the derived slope (0.416) and intercept (−0.155 kcal/mol) values to recenter the energy terms of the combinatorial mutants.

### Visualisation of numeric values on Aβ42 structures

For purposes of visualisation of numeric values (energy terms, ΔΔE_a_/ΔΔG ratios and energetic couplings) on Aβ42 structures in figures, we used the following chains for each of the PDB files: B for 2BEG; D for 2MXU; C for 2NAO; D for 5KK3; F for 5OQV; E for 7Q4B; G for 7Q4M; E for 8EZD; and E for 8EZE. Molecular graphics and analyses performed with UCSF ChimeraX, developed by the Resource for Biocomputing, Visualization, and Informatics at the University of California, San Francisco, with support from National Institutes of Health R01-GM129325 and the Office of Cyber Infrastructure and Computational Biology, National Institute of Allergy and Infectious Diseases^66^.

### Fibril stability analyses

Inspired by phi-value analysis^17,24^, we calculated activation energy to fibril stability energy ratios for Aβ42 by dividing our inferred activation energy terms ΔΔE_a_ (Fig. 2f) by the change in free energy ΔΔG of fibril state structures^46^ of Aβ42 (separately for 2BEG, 2MXU, 2NAO, 5KK3, 5OQV, 7Q4B, 7Q4M, 8EZD, 8EZE, see Supplementary Table 5 for structure details). Following the example of a recent study on PI3K-SH3 amyloids, where FoldX predictions of ΔΔG were shown to correlate well with *in vitro* measurements of fibrils stability for 15 PI3K-SH3 variants^24^, we ran FoldX on a stacked single filament tetramer fibril structures of Aβ42 peptides. For the 2NAO structure we ran it on a stacked single filament trimer conformation, as the PDB structure did not contain more than three stacked chains in any filament. We used the following Aβ42 chains: B, C, D, E for 2BEG; A, B, C for 2NAO; A, B, C, D for 2MXU; A, B, C, D for 5KK3; A, C, F, H for 5OQV; B, D, F, R for 7Q4B; A, C, E, G for 7Q4M; E, F, G, H for 8EZD; and E, F, G, H for 8EZE. Resulting ΔΔG values were then divided by 4 (or 3 in case of 2NAO) to obtain per-monomer ΔΔG values. In our downstream analysis, we further only considered those ΔΔE_a_/ΔΔG ratios, for which the predicted ΔΔG values were above 0.6 kcal/mol (literature-motivated threshold^23^) and below 10 kcal/mol, to only consider mutations that do not significantly perturb FS structure (Fig. 3b,c, Fig. S4, Fig. S5a-h). The main assumption underlying ΔΔE_a_/ΔΔG ratio analysis is that the introduced mutations do not perturb these structures drastically.

### Calculation of inter-residue distances in 3D space

Distances between AA side chains in 3D space were calculated using the DMS2structure toolkit (available at https://github.com/lehner-lab/DMS2structure/ and published in a previous study^52^). Briefly, for two given amino acids in a PDB structure, DMS2structure calculates distances between all pairs of heavy atoms (any atoms other than hydrogen) in their side chains across the two amino acids. The minimum of these distances is then reported as the minimal side chain heavy atom distance (scHA_min), further used in downstream analysis. In absence of a side chain for Glycine, the central carbon atom (C-α) is used for calculations. We specifically used *contact_matrix_from_pairdistances.R* script to calculate inter-residue distances for monomer conformations of Aβ42 structures, and *pairdistances_from_PDB_crystal.R* script to calculate inter-residue distances for dimer (2 monomers in different filaments facing each other) conformations of Aβ42 structures.

## Data availability

Raw sequencing data and the processed data table (Supplementary Table 2) are deposited in NCBI’s Gene Expression Omnibus (GEO) with accession number GSE269461 (N-terminal Aβ42 double mutant, C-terminal Aβ42 double mutant and combinatorial libraries) and GSE151147 (shallow double mutants library^26^).

## Code availability

All the code used for analyses presented in this work are available at: https://github.com/lehner-lab/amyloids_energy_modelling.

## Author contributions

A.A. and M.S. analysed the data. M.S. performed all experiments. M.S., B.B. and B.L. designed the experiments. A.J.F. developed the modelling approach and performed preliminary data analyses with M.S.. B.B. and B.L. conceived the project and supervised the research. A.A., B.B. and B.L. wrote the manuscript with input from M.S..

## Acknowledgements

This work was funded by the La Caixa Research Foundation project ‘DeepAmyloids’ (LCF/PR/HR21/52410004). Work in the lab of BB was also funded by the Spanish Ministry of Science, Innovation and Universities (PID2021-127761OB-I00, RYC2020-028861-I funded by MCIN/AEI/ 10.13039/501100011033, “ERDF A way of making Europe” and “ESF Investing in your future”) and by the European Union (ERC Consolidator, Glam-MAP, 101125484). Views and opinions expressed are however those of the author(s) only and do not necessarily reflect those of the European Union or the European Research Council. Neither the European Union nor the granting authority can be held responsible for them. Work in the lab of B.L. was funded by a European Research Council (ERC) Advanced (883742) grant, the Spanish Ministry of Science and Innovation (LCF/PR/HR21/52410004, EMBL Partnership, Severo Ochoa Centre of Excellence), the Bettencourt Schueller Foundation, the AXA Research Fund, Agència de Gestió d’Ajuts Universitaris i de Recerca (AGAUR, 2017 SGR 1322), the CERCA Program/Generalitat de Catalunya and Wellcome (Grant reference: 220540/Z/20/A, ’Wellcome Sanger Institute Quinquennial Review 2021-2026’). M.S was funded by a fellowship from Agencia de Gestio d’Ajuts Universitaris i de Recerca (2019FI_B 01311). A.J.F. was funded by a Ramón y Cajal fellowship (RYC2021-033375-I) financed by the Spanish Ministry of Science and Innovation (MCIN/AEI/10.13039/501100011033) and the European Union (NextGenerationEU/PRTR).

## Description of Supplementary Tables

**Supplementary Table 1**

Description: Primary and secondary nucleation rate constants measured independently in previous studies^36,37^ for Aβ42 mutants (with each of the studies’ data in separate sheets), and their corresponding relative growth rates measured in this study in the double mutants datasets (GR_shallow_double_mutants, GR_C_terminal_Ab42_double_mutants and GR_N_terminal_Ab42_double_mutants) and change in additive trait as inferred by MoCHI^34,35^ (ddEa_joint_model) trained on all double mutants datasets. For the Yang et al. dataset^37^ primary and secondary nucleation rate constants (k_n_ and k_2_) reported in the study are in ‘kn_yang’ and ‘k2_yang’ columns, respectively. For the Thacker^36^ dataset, multiplicative terms k_+_k_n_ and k_+_k_2_ (primary and secondary nucleation rate constants multiplied by the rate of elongation) were derived from the reported λ and κ values (the rate at which new fibril mass is formed via primary and secondary nucleation, respectively, in columns ‘lambda’ and ‘kappa’) using the exact model description authors provided in the supplementary material of the publications, and are in ‘k+kn_thacker’ and ‘k+k2_thacker’ columns, respectively.

**Supplementary Table 2**

Description: Relative growth rates (relative to WT) and their associated error estimates from DiMSum^64^ for all the Aβ42 mutants (double and combinatorial) generated and analysed in this study.

**Supplementary Table 3**

Description: Inferred changes in activation energy (column ‘ddEa_scaled’, in kcal/mol units) for all the substitutions in Aβ42 (column ‘id’) with associated outputs from MoCHI^34,35^ model (fit on all the Aβ42 double mutant datasets jointly). Columns ‘zscore_unaffected’, ‘p.adjust_mode’, ‘category_affected’ and ‘category_incr_decr_nucleation’ present the statistics from the Z-test asking whether inferred ΔΔE_a_ values are different from ΔΔE_a_ = 0 kcal/mol (unaffected nucleation) and in which direction (> 0 or < 0). For the first entry (WT) ΔE_a_ is reported.

**Supplementary Table 4**

Description: (ddEa_and_GR_values sheet) inferred changes in activation energy (column ‘ddEa’), corresponding mean relative growth rate for all Aβ42 double mutants (for each mutant, the mean relative growth rate is calculated across all three or less Aβ42 double mutants datasets where this mutant is present, with column ‘dataset’ indicating which dataset the mutant is present in). (ddEa_and_GR_Zscores sheet) Z-scores of changes in activation energy (X column) and Z-scores of relative growth rates necessary to reproduce heatmaps in ED Fig.3b-d (columns ‘ddEa_norm’, ‘neg_GR_mean_norm’ and ‘ddEa_norm_minus_neg_GR_mean_norm’, respectively, for panels b,c and d).

**Supplementary Table 5**

Description: Aβ42 structures used in analyses of the study, indicating their PDB ID, technique employed for structural determination and peptide origin. H-D - Hydrogen Deuterium, NMR - Nuclear Magnetic Resonance, SS - solid state, EM - electron microscopy, Cryo EM - cryogenic electron microscopy, MAS NMR - magic-angle spinning NMR.

**Supplementary Table 6**

Description: Processed FoldX output table containing predicted changes in free energy (ΔΔG) of Aβ42 fibril structures (each sheet in the table corresponds to a given Aβ42 PDB structure). Column ‘total energy’ contains total ΔΔG predicted for a tetramer (or in case of 2NAO structure - trimer, see Methods for details) of Aβ42 fibrils, whereas column ‘ddG_per_monomer’ contains ΔΔG values predicted by FoldX for a single monomer fibril of Aβ42. All energies are in kcal/mol units.

**Supplementary Table 7**

Description: Ratios of activation energy change ΔΔE_a_ to fibril stability change ΔΔG (column ‘ddEa_to_ddG_ratio’) for all the relevant substitutions in Aβ42 (those with ΔΔG satisfying the following condition: 0.6 kcal/mol < |ΔΔG| < 10 kcal/mol, see Methods for more details); corresponding changes in activation energy (column ‘ddEa’) and free energy of fibril state (column ‘ddG’) that were used to calculate the phi values, for all available Aβ42 structures (separately in each sheet of the table). All energies are in kcal/mol units.

**Supplementary Table 8**

Description: Root mean square distance to 1 across all ΔΔE_a_ /ΔΔG ratios (R) in APR2 region of Aβ42 (AA 29-42) (‘rms_indiv_ddEa_to_ddG_ratios’ column) for each PDB structure of Aβ42 used in the analysis.

**Supplementary Table 9**

Description: Inferred changes in activation energy (‘ddEa’ sheet, column ‘ddEa_scaled’, in kcal/mol units) and energetic couplings (‘dddEa’ sheet, column ‘dddEa_scaled’, in kcal/mol units) with associated outputs from MoCHI^34,35^ model (fit on all the Aβ42 combinatorial mutant datasets jointly) for all the mutations and their pairwise combinations introduced in the combinatorial Aβ42 mutant datasets.

**Supplementary Table 10**

Description: Interaction scores (calculated as mean of absolute energetic couplings, ‘mean_abs_dddEa’ column) and the corresponding 3D distance between residues (‘scHA_min’ column) for every pair of positions (‘Ab42_position_pair’ column) mutagenised in Aβ42 combinatorial mutant datasets (these data are used to produce scatterplots in Fig. 4e and ED Fig. 9a).

**Supplementary Table 11**

Description: Complete list of oligonucleotides employed for cloning, mutagenesis and sequencing library preparation in this study.

## Supplementary Figures

**Figure S1.**
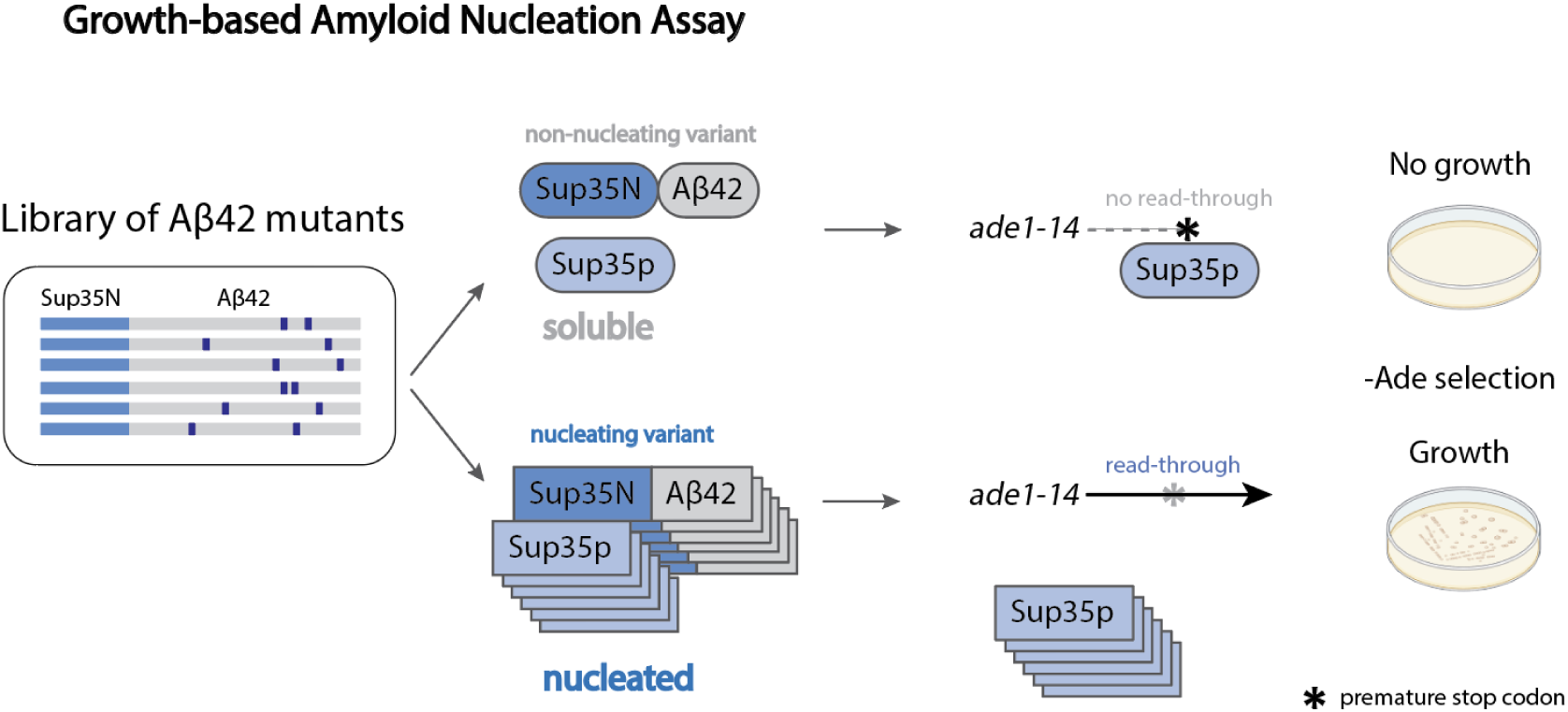
High throughput quantification of amyloid nucleation kinetics. Schematic overview of the amyloid nucleation assay: Aβ42, fused to the nucleation domain of Sup35 (Sup35N), seeds aggregation of the yeast prion Sup35p, causing read-through of a premature stop codon in the *ade1* reporter gene, thus allowing growth in medium lacking adenine.

**Figure S2.**
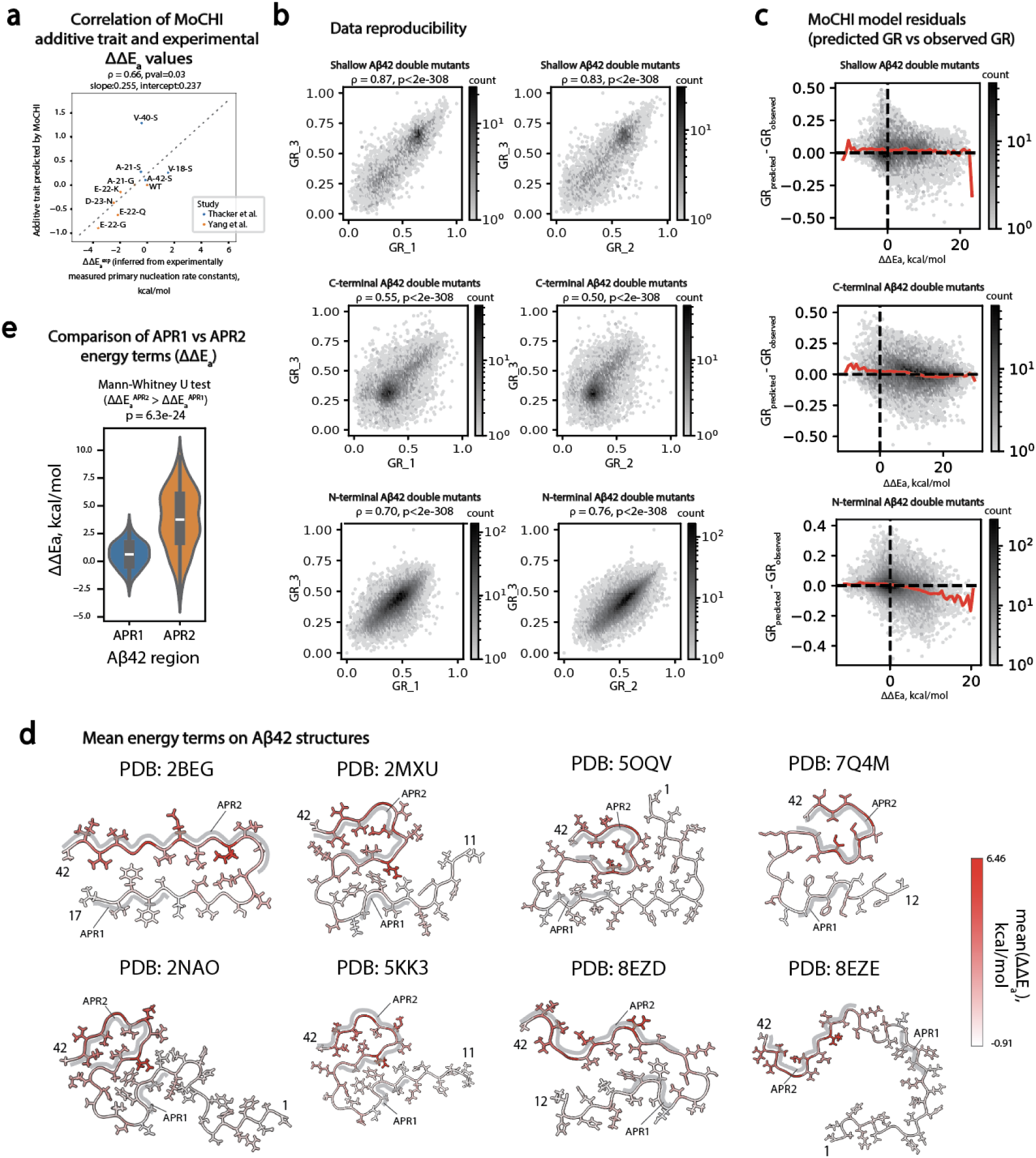
Overview of double mutant libraries and inferred activation energy changes. **a**, Scatterplot of additive trait values predicted by MoCHI^35^ model trained on double mutant datasets (Y axis) and experimentally-derived ΔΔE_a_ values (X axis, using primary nucleation rate constants). Dashed grey line represents linear regression fit for the data. **b**, Inter-replicate correlations of relative growth rates (GR) for shallow, C-terminal and N-terminal Aβ42 double mutant libraries (from top to bottom). Spearman’s ρ (correlation) coefficients and associated p-values are reported. **c**, MoCHI residuals (predicted vs observed relative growth rates, red line is following mean residuals in 50 equally spaced bins across x axis, dashed black lines indicate 0 in both axes) for shallow, C-terminal and N-terminal Aβ42 double mutant libraries (from top to bottom). **d** Cross sections of 2BEG, 2MXU, 5OQV, 7Q4B, 2NAO, 5KK3, 8EZD and 8EZE PDB structures of Aβ42 fibrils coloured by mean ΔΔE_a_ per position. Aggregation prone regions 1 (APR1) and 2 (APR2) are highlighted in grey. **e**, Violinplot comparing inferred activation energy terms (ΔΔE_a_) between APR1 region (AA 17-21) and APR2 region (AA 29-42) of Aβ42 (p=6.3e-24, one-sided Mann-Whitney U test (ΔΔE ^APR^^2^ > ΔΔE ^APR^^1^)).

**Figure S3.**
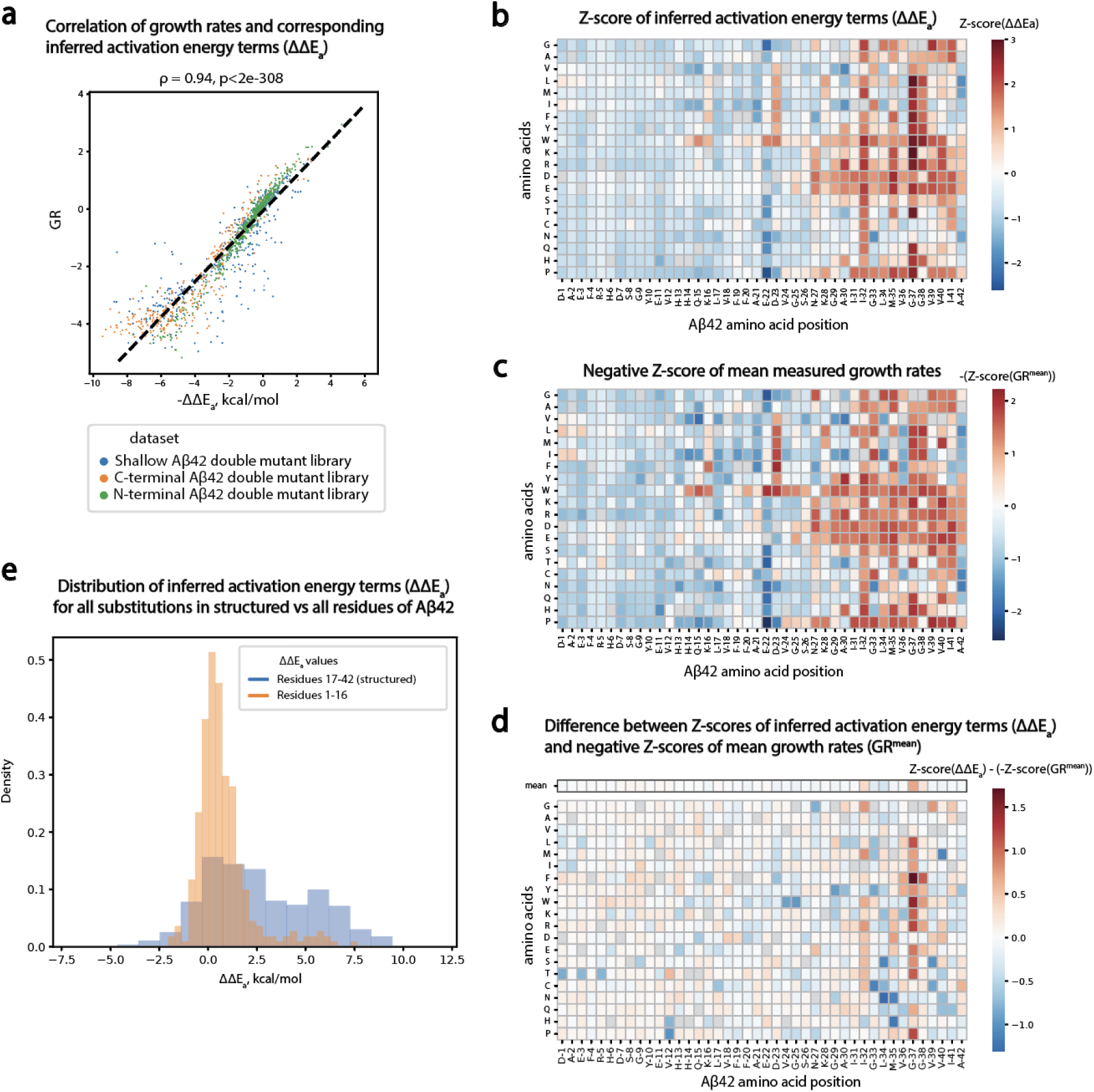
Comparison of activation energies and cellular relative growth rates. **a**, Correlation of relative growth rates of shallow, C-terminal and N-terminal Aβ42 double mutants and corresponding inferred activation energy terms (ΔΔE_a_). Spearman’s ρ (correlation) coefficients and associated p-values are reported. Dashed black line represents linear regression fit to the data. **b**, Z-scores of inferred activation energy terms (ΔΔE_a_) (scaled to zero mean and unit variance). **c**, Negative Z-scores of mean (across all double mutant datasets) relative growth rates (GR^mean^) (scaled to zero mean and unit variance). **d**, Difference between Z-scores of inferred activation energy terms (ΔΔE_a_) and negative Z-scores of mean relative growth rates (GR^mean^). Means of difference values for each position are displayed in the top row of the heatmap (outlined in black). e, Distribution of inferred activation energy terms (ΔΔE_a_) for all substitutions across only those Aβ42 positions that are structurally resolved in all available fibril structures (AA 17-42, in blue) or the rest of the positions in Aβ42 (AA 1-16, in orange).

**Figure S4.**
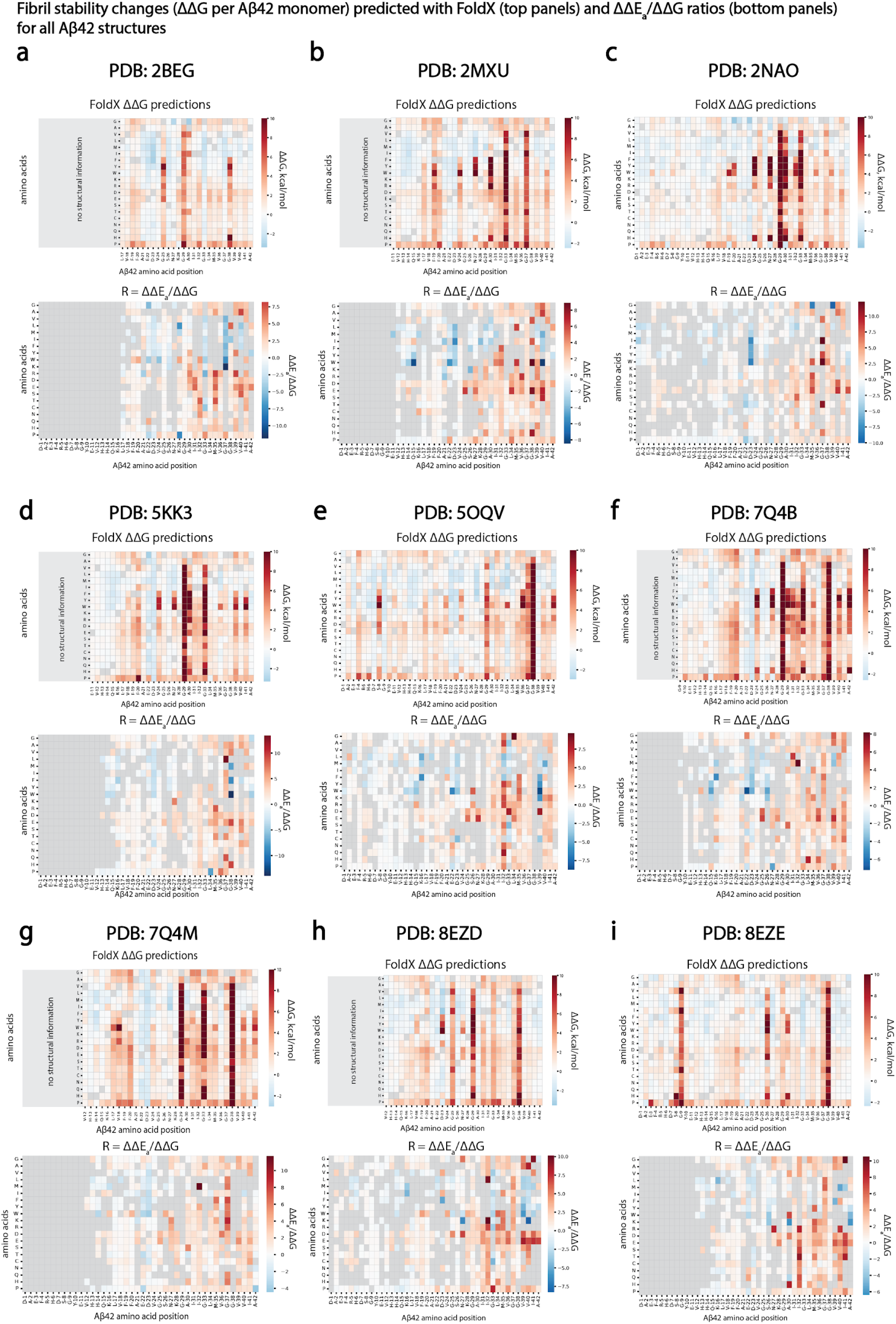
Fibril stability and ΔΔE_a_/ΔΔG ratios (R) for all Aβ42 structures. **a-i**, (top panels) Fibril stability changes (ΔΔG per Aβ42 monomer) predicted with FoldX upon all possible substitutions and (bottom panels) R-values for structures 2BEG (a), 2MXU (b), 2NAO (c), 5KK3 (d), 5OQV (e), 7Q4B (f), 7Q4M; (g) 8EZD (h), 8EZE (i). FoldX was run using a stack of four Aβ42 monomers for all structures, apart from 2NAO where a stacked trimer was used (see Methods).

**Figure S5.**
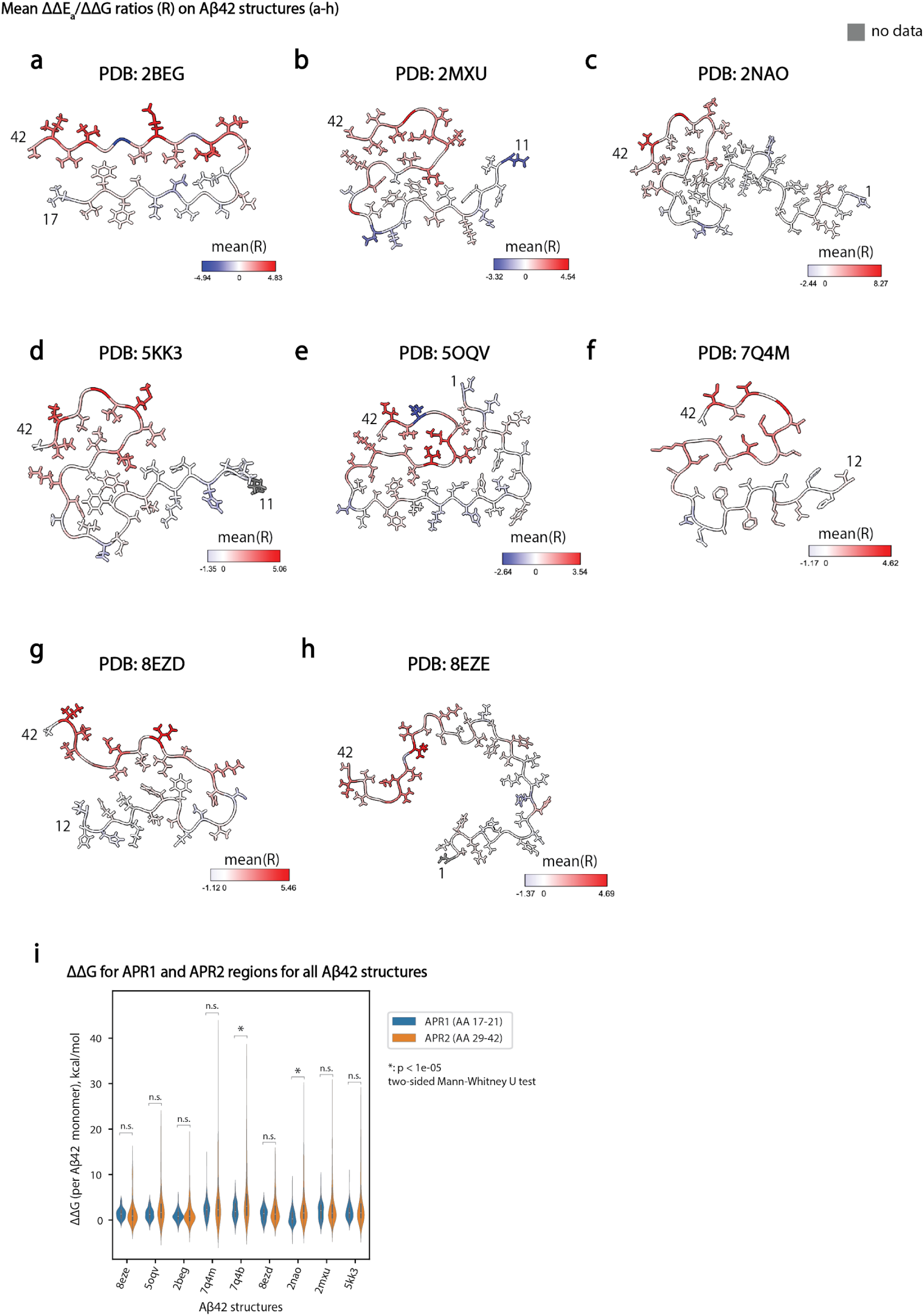
Mean ΔΔE_a_/ΔΔG ratios (R) on all Aβ42 structures. **a-h**, Cross sections of PDB structures of Aβ42 fibrils coloured by mean R-values at that position: 2BEG (a), 2MXU (b), 2NAO (c), 5KK3 (d), 5OQV (e), 7Q4B (f), 8EZD (g), 8EZE (h). Residues with no R-values are coloured in grey. i, Violin plot comparing fibril stability changes (ΔΔG per Aβ42 monomer predicted with FoldX) for all possible substitutions in APR1 vs APR2 regions of Aβ42, for all available Aβ42 fibril structures (one-sided Mann-Whitney U test statistics are reported).

**Figure S6.**
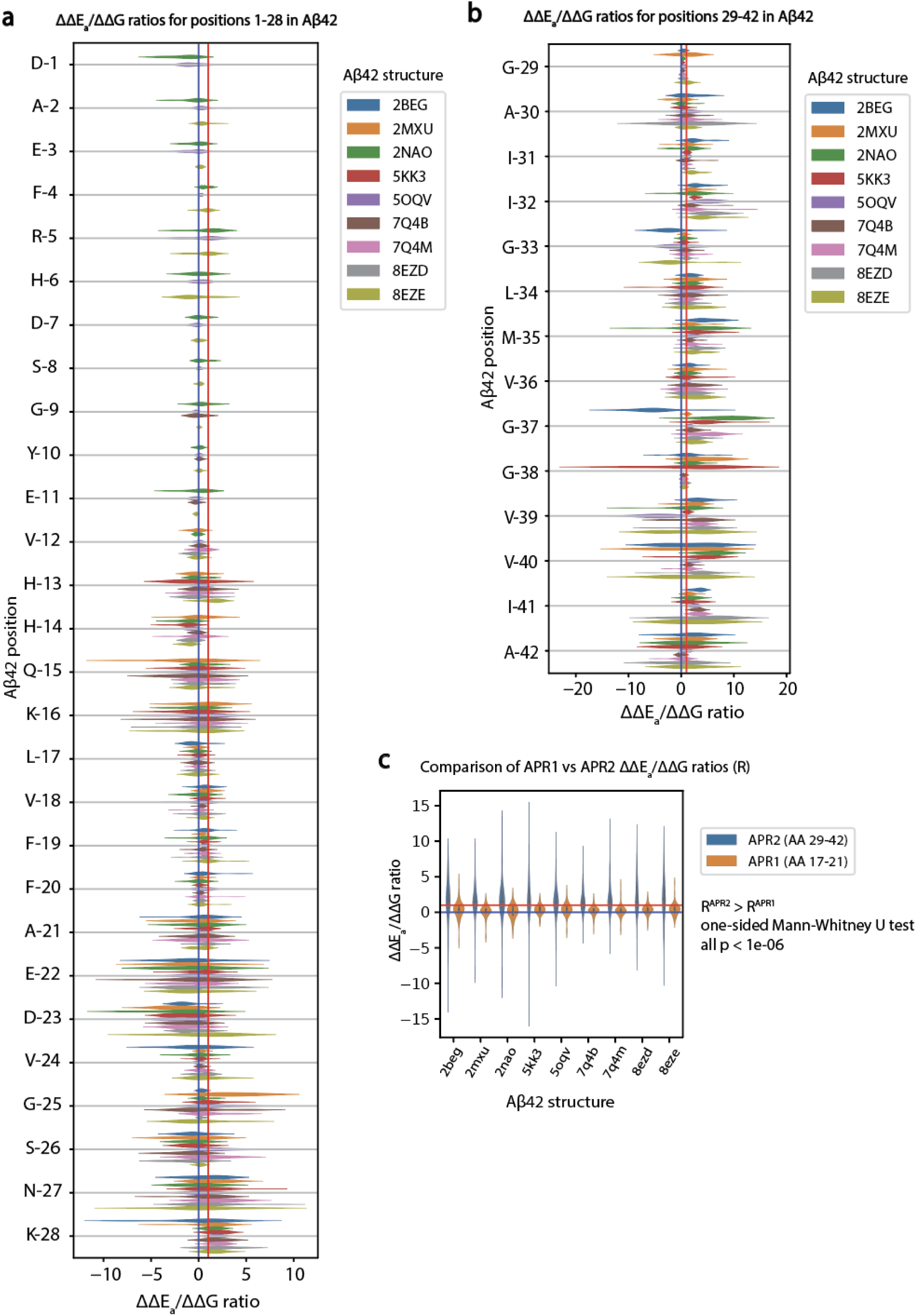
Overview of all R-values and comparison of APR1 and APR2 R-values. **a-b**, Violinplot of R-values for all the Aβ42 structures (2BEG, 2MXU, 2NAO, 5KK3, 5OQV, 7Q4B, 7Q4M, 8EZD, 8EZE) for each position at the (a) N-terminus (AA 1-27) and (b) C-terminus (AA 28-42). Red vertical lines mark R = 1, and blue vertical lines mark R = 0. **c**, Violinplot comparing APR1 region (AA 17-21) and APR2 region (AA 29-42) R-values for all the Aβ42 structures (2BEG, 2MXU, 2NAO, 5KK3, 5OQV, 7Q4B, 7Q4M, 8EZD, 8EZE). One-sided Mann-Whitney U test statistics are reported (p<1e-06 for all comparisons).

**Figure S7.**
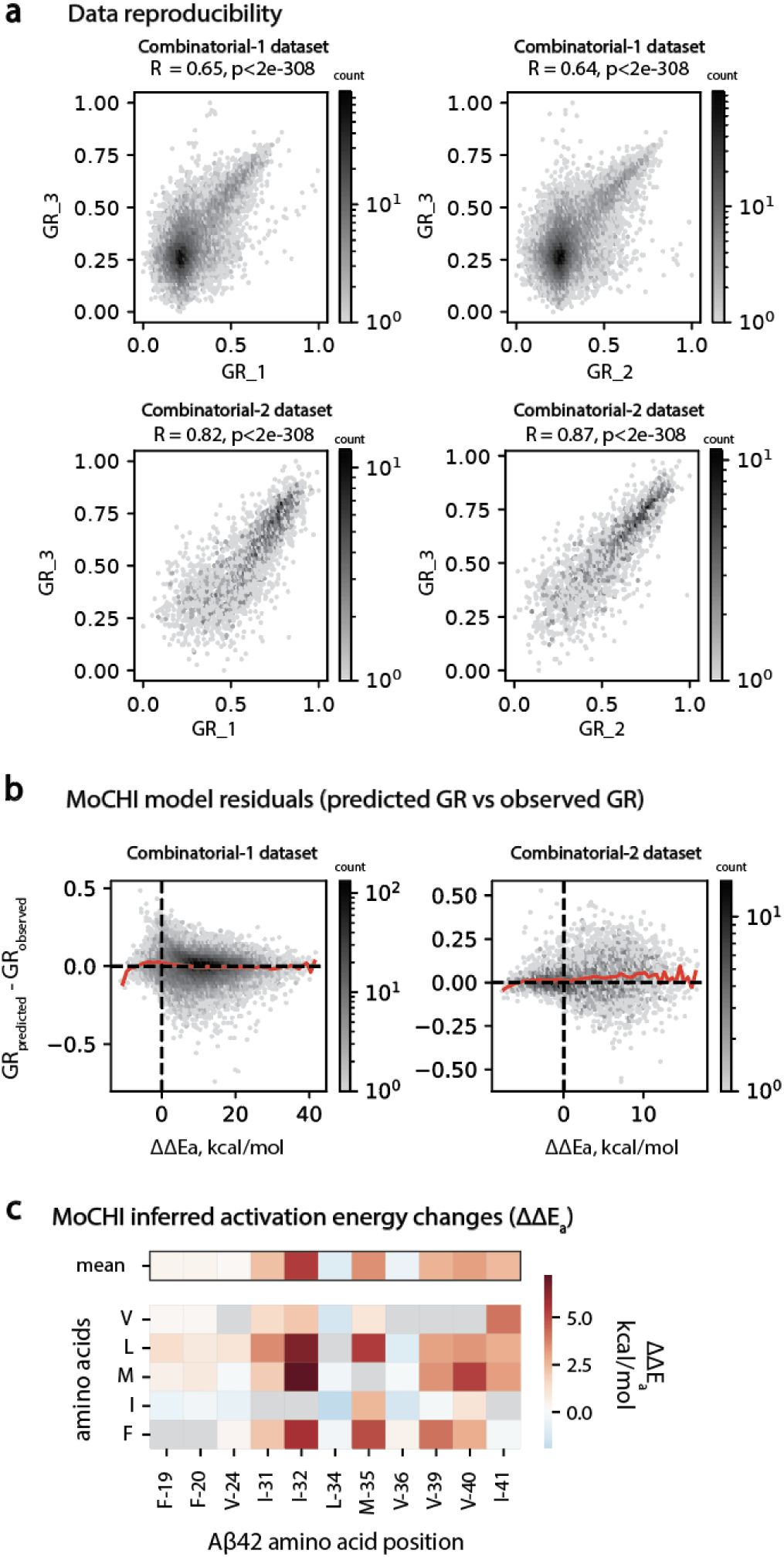
Overview of combinatorial mutant libraries and energetic couplings. **a**, Inter-replicate correlations of relative growth rates for Combinatorial-1 (top panels) and Combinatorial-2 (bottom panels) combinatorial mutant libraries. Pearson’s correlation coefficients (R) and associated p-values are indicated. **b**, MoCHI residuals (predicted vs observed relative growth rates, red line is following mean residuals in 50 equally spaced bins across x axis, dashed black lines indicate 0 in both axes) for Combinatorial-1 (left) and Combinatorial-2 (right) combinatorial mutant libraries. **c**, Heatmap displaying the inferred activation energy changes (ΔΔE_a_) for all Aβ42 substitutions present in combinatorial mutagenesis datasets (Combinatorial-1, Combinatorial-2). Mean ΔΔE_a_ values for each position are displayed in the top row of the heatmap (outlined in black).

**Figure S8.**
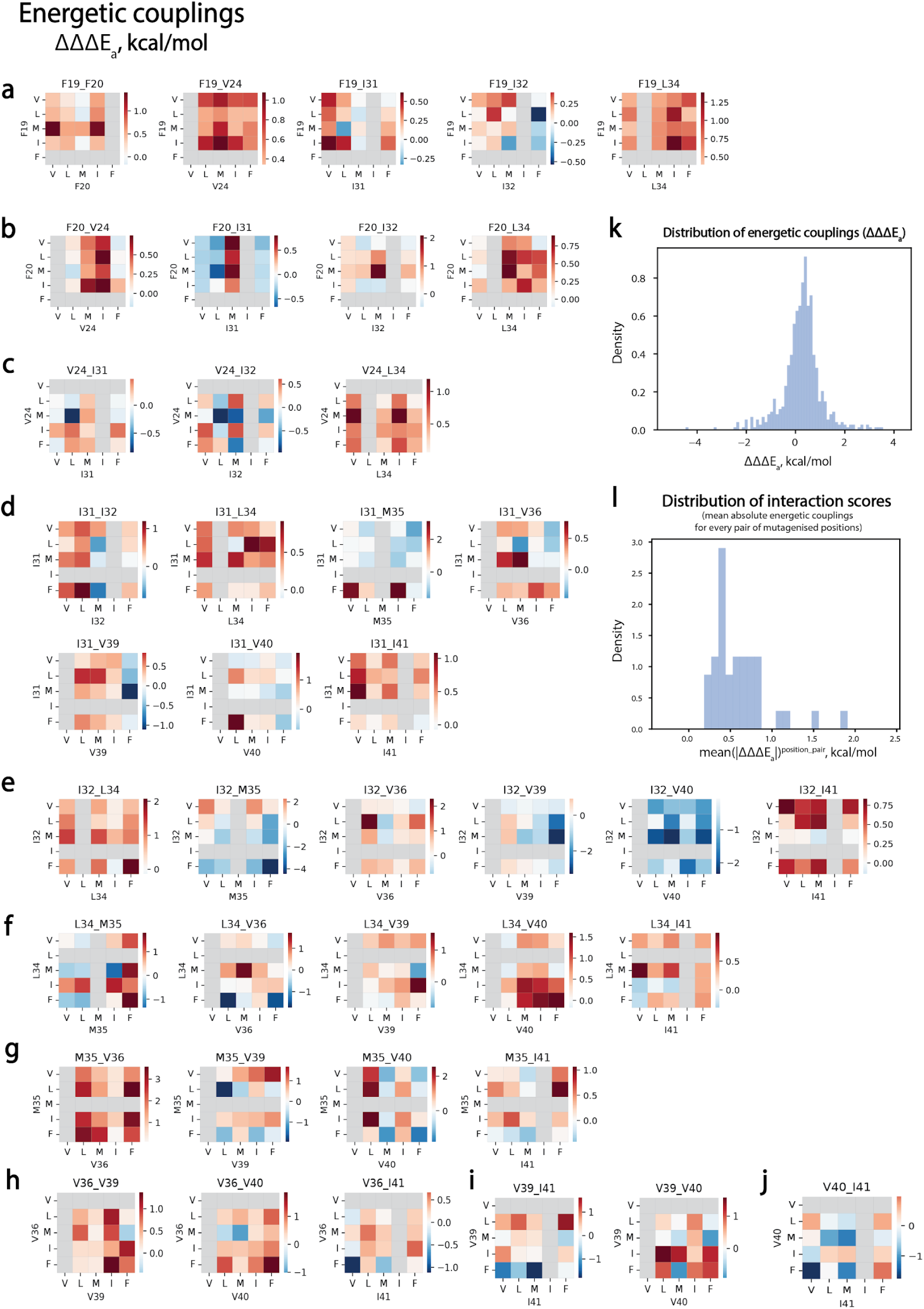
Energetic coupling landscape of Aβ42. **a-j**, Heatmaps displaying the energetic couplings between all individual mutations introduced in the combinatorial mutagenesis datasets (Combinatorial-1, Combinatorial-2) for each pair of mutated positions (F-19 (a), F-20 (b), V-24 (c), I-31(d), I-32 (e), L-34 (f), M-35 (g), V-36 (h), V-39 (i), V-40 (j)). **k**, Distribution of 640 energetic couplings (ΔΔΔE_a_) between mutations introduced in combinational Aβ42 mutant datasets. l, Distribution of 40 interaction scores (mean |ΔΔΔE_a_|) for pairs of positions mutagenised in combinatorial Aβ42 mutant datasets.

**Figure S9.**
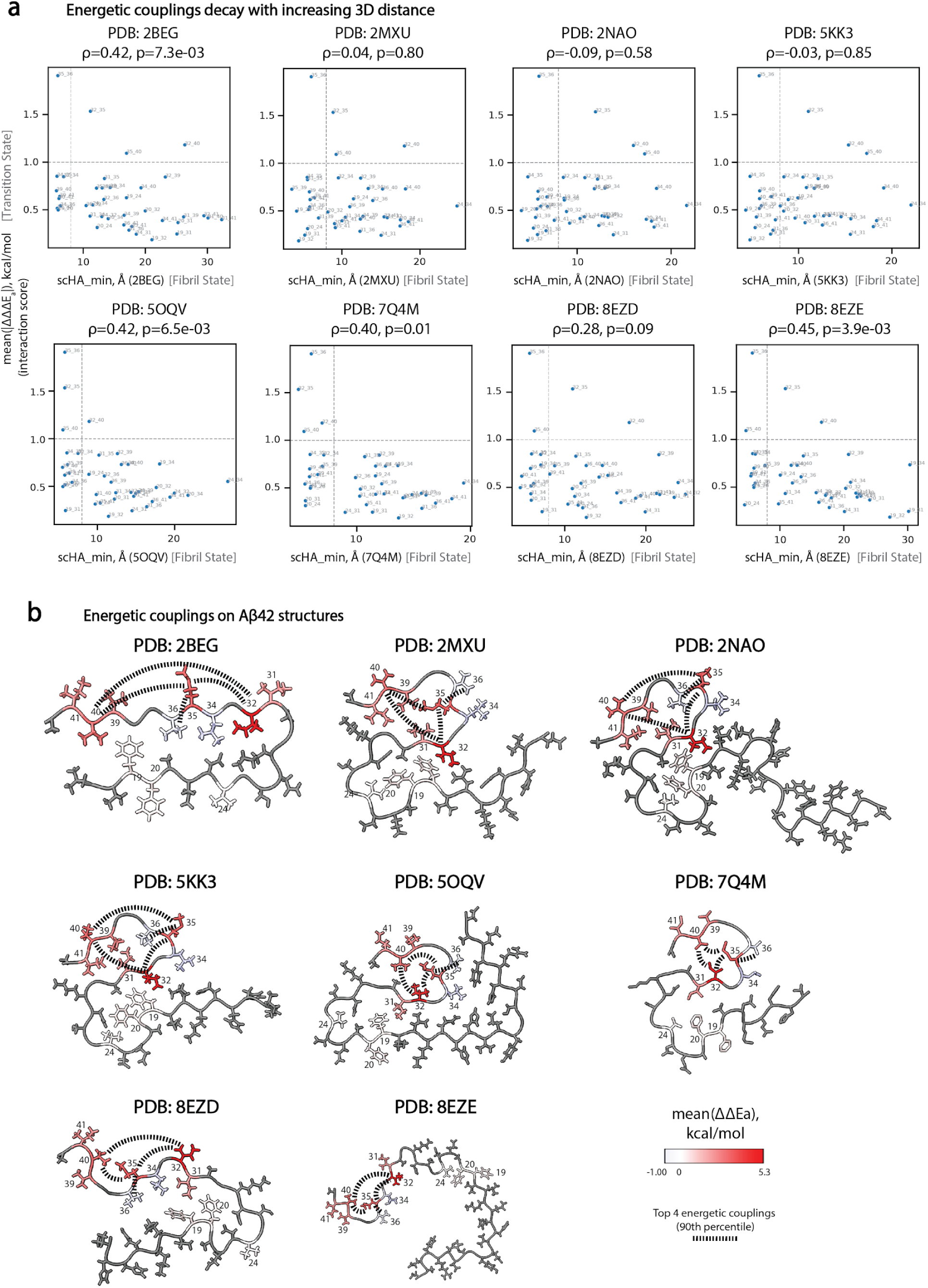
Energetic couplings compared to Aβ42 fibril polymorphs. **a**, Scatterplots of interaction scores for pairs of positions and the inter-residue distance for corresponding pairs of amino acids in 3D space (scHA_min, minimum heavy atom side chain distance) of Aβ42 structures 2BEG, 2MXU, 2NAO, 5KK3, 5OQV, 7Q4M, 8EZD, 8EZE; dashed light grey vertical line marks 8 Å, dashed light grey horizontal line marks interaction score (mean(|ΔΔΔE_a_|) of 1 kcal/mol. **b**, Cross sections of PDB structures (2BEG, 2MXU, 2NAO, 5KK3, 5OQV, 7Q4M, 8EZD, 8EZE) of Aβ42 fibrils coloured by mean inferred activation energy terms (ΔΔEa) from the MoCHI model trained on combinatorial mutants datasets (residues with no inferred ΔΔEa values are in grey). Positions mutagenised in combinatorial datasets (Combinatorial-1 and Combinatorial-2) are labelled on the PDB structure. Top 4 interacting position pairs (in 90^th^ percentile by their interaction scores) are connected with dashed black lines.

